# Differential Allele-Specific Expression Uncovers Breast Cancer Genes Dysregulated By *Cis* Noncoding Mutations

**DOI:** 10.1101/675462

**Authors:** Pawel F Przytycki, Mona Singh

## Abstract

Identifying cancer-relevant mutations in noncoding regions is extremely challenging due to the large numbers of such mutations, their low levels of recurrence, and the general difficulty in interpreting their impact. To uncover genes that are dysregulated due to somatic mutations in *cis*, we build upon the concept of *differential allele-specific expression* (ASE) and introduce methods to identify genes within an individual’s cancer whose ASE differs from what is found in matched normal tissue. When applied to breast cancer tumor samples, our methods readily detect the known allele-specific effects of copy number variation and nonsense-mediated decay. Further, genes that are found to recurrently exhibit differential ASE across samples are cancer relevant. Genes with *cis* mutations are enriched for differential ASE, and we find 147 potentially functional noncoding mutations *cis* to genes that exhibit significant differential ASE. Overall, our results suggest that differential ASE is a promising means for discovering gene dysregulation within an individual due to *cis* noncoding mutations.

## Introduction

Cancer is a heterogeneous disease in which every tumor exhibits a unique combination of genotypic and phenotypic features (Garraway & Lander, 2013). A major aim of cancer genomics is to understand how acquired somatic mutations within cancer cells lead to the dysregulation that allows cells to grow and proliferate uncontrollably (Vogelstein et al., 2013). Towards this aim, consortia such as TCGA (TCGA Research Network) and ICGC (International Cancer Genome Consortium, 2010) have profiled the tumors of thousands of individuals across tens of cancer types to determine DNA sequences, gene expression levels, copy number variation, and other functional genomics data. Initial analyses have revealed that each individual’s tumor typically has a unique combination of mutations, most of which play no functional role in cancer (Garraway & Lander, 2013; Vogelstein et al., 2013).

While many computational methods have been developed to identify which of the numerous somatic alterations observed within a tumor are linked to cancer phenotypes, these have mostly focused on mutations affecting protein-coding regions (Diederichs et al., 2016). However, the vast majority of cancer somatic mutations are found in noncoding regions (Khurana et al., 2016), as ~98% of the human genome is noncoding (Venter et al., 2001; Lander et al., 2001). Further, since widespread gene expression dysregulation is common in cancer and the regulatory elements controlling gene expression are largely found within the noncoding portion of the genome (The ENCODE Project Consortium, 2012), methods to link noncoding somatic alterations within an individual’s tumor to changes in gene expression would not only vastly advance our knowledge of cancer biology but also be a great aid in interpreting specific cancer genomes.

Perhaps the most commonly applied approaches to identify functional somatic mutations in cancer search for mutations that recur across tumor samples more than expected by chance. In the context of noncoding mutations, such approaches may additionally integrate functional impact scores or diverse functional genomics annotations relevant for gene regulation (Fu et al., 2014; Lochovsky et al., 2015; Mularoni et al., 2016). Overall, while recurrence-based approaches have proven to be highly successful in identifying cancer-relevant mutations within coding regions (Lawrence et al., 2013; Przytycki & Singh, 2017), they have had limited success in uncovering such mutations within noncoding regions (Khurana et al., 2016). The best understood of these are mutations within the *TERT* promoter, which occur in about 30% of melanomas and are relatively common in other cancers and are associated with *TERT* overexpression (Huang et al., 2013; Vinagre et al., 2013). However, further recurrence analysis across hundreds of deeply sequenced samples has only uncovered a handful of other candidate causal mutations in cancer (Melton et al., 2015; Nik-Zainal et al., 2016; Rheinbay et al., 2017; Weinhold et al., 2014).

A parallel approach is to link somatic noncoding mutations to changes in gene expression. Samples with and without mutations within a proximal upstream region of a specific gene can be tested for association with different expression levels for that gene (Fredriksson et al., 2014; Gyorffy et al., 2018). Mutation and gene pairs can also be evaluated in the context of known functional enhancers and promoters, thereby allowing the inference of more distally mutated loci associated with changes in expression in cancers (Feigin et al., 2017; Hornshøj et al., 2018; W. Zhang et al., 2018). However, in order to infer statistically significant differences in gene expression, these types of approaches also require that noncoding regulatory regions for a gene are recurrently mutated across individuals.

Our work begins with the observation that functional noncoding somatic mutations within regulatory regions should change the expression of one copy of the target gene without affecting the expression of the allele on the other chromosome. Thus, we aim to identify genes where only the expression of one allele is changed in cancer cells in order to hone in on *cis* noncoding mutations that lead to regulatory changes that are functional in cancers. However, unequal expression levels of alleles of a gene, or allele-specific expression (ASE), is a relatively common phenomenon in both healthy tissues and cancer (Aguet et al., 2017; Mayba et al., 2014; Romanel et al., 2015; Spurr et al., 2018). While ASE has been used to measure the impact of cis-eQTLs across healthy tissues (Mohammadi et al., 2017), in the context of cancer, ASE is often detected in more than 25% of all genes (Ma et al., 2018; Mayba et al., 2014; Romanel et al., 2015), making it difficult to apply for detecting either genes dysregulated in cancers due to alterations in *cis* or underlying functional noncoding somatic mutations.

To help narrow the search for genes whose regulation is affected by changes in *cis*, we introduce new methods to detect *differential ASE* in the context of cancer, where the ASE of a gene within cancer cells is different from its ASE in paired normal cells from the same individual and tissue type. Differential ASE aims to find dysregulated genes within an individual by directly comparing the level of ASE in a tumor sample to the level of ASE at the same heterozygous site in a matched normal sample. In this way, the only genomic sequence differences between the paired samples are cancer related, and thus changes in ASE are as well. However, detecting ASE in tumor cells is complicated by intra-tumor heterogeneity. We thus introduce a series of increasingly complex models to detect differential ASE from gene expression data, including two models to deconvolve expression measurements derived from tumor samples in order to estimate the contributions from cancer cells.

Most previous studies of ASE in cancer have not directly compared ASE levels for genes in tumors with their matched normal counterparts. For example, while genes with different levels of ASE have been observed when comparing cell lines derived from the germlines of those with familial pancreatic cancer with control samples, ASE in tumor cells was not considered (Aik et al., 2008). An alternate approach which considered somatic coding mutations and uncovered those mutations that showed preferential ASE in tumor samples did not consider whether genes already exhibited ASE in that individual’s normal tissue (Halabi et al., 2016). Another approach has been to test whether genes that have significant ASE in cancer tissues also have significant ASE in matched normal tissues in order to classify the effect as tumor-specific or shared (Ongen et al., 2014). However, this merely determines if a gene as a whole exhibits significant ASE in one context but not the other.

Previous usage of differential ASE in the cancer context has been limited. For example, ASE and differential ASE have been used to detect copy number variation in tumor samples (e.g., Attiyeh et al., 2009; Peiffer et al., 2006). Further, the software MBASED for detecting ASE includes a two-sample version, which can be used to determine if there is a significant difference in a gene’s level of ASE in a tumor versus normal sample (Mayba et al., 2014). Our differential ASE approach differs from these previous works as it is, to the best of our knowledge, the first aimed at detecting the effects of *cis* noncoding somatic mutations while also being able to account for tumor heterogeneity and different levels of expression between paired tumor and normal samples.

We apply our approach to detect genes that exhibit differential ASE to paired breast cancer and normal samples profiled by TCGA (TCGA Research Network). First, as validation, we demonstrate that the dysregulation caused by copy number variation and nonsense-mediated decay results in differential ASE. Next, we observe that genes that recurrently exhibit differential ASE include known cancer drivers and fall in cancer-related pathways. Furthermore, estrogen receptor positive tumors tend to show differential ASE of estrogen receptor genes. We then focus in on noncoding mutations derived from whole-exome sequencing data, as very few of the individuals with paired expression profiles have whole-genome sequencing data. We find that genes with *cis* noncoding mutations are enriched for differential ASE. Of the observed *cis* noncoding mutations, 96.2% do not result in differential ASE of the corresponding gene, and thus can be filtered out as they are highly unlikely to be functional. Of the remaining mutations, we find 147 mutations that occur in annotated regulatory sites. We extend this analysis by looking at whole-genome sequencing data in four individuals and identify an additional five mutations within annotated regulatory sites that are upstream of genes that exhibit differential ASE. We conclude that differential ASE is a powerful approach to hone in on functional *cis* noncoding mutations.

## Results

### Overview of Differential Allele-Specific Expression

We define the differential ASE of a gene as the difference between its estimated ASE in cancer cells and a paired normal tissue sample. Briefly, rather than computing ASE at each heterozygous site by comparing the fraction of RNA-seq reads in tumor samples that contain a specific nucleotide to an expected fraction of one half (as is done to estimate ASE when using only a single sample), we consider the observed fraction at the same site in a matched normal sample. Because tumor samples are heterogeneous and contain both cancer and normal cells, estimating ASE in cancer cells is challenging and we introduce three models for this task (see Figure 1). In the first model, diffASE-baseline, we directly compute the difference between the fraction of transcripts with the specific nucleotide in the tumor sample and the normal sample. In the second model, diffASE-purity, we use estimated tumor purity to deconvolve expression data from the tumor sample into proportions arising from cancer cells and normal cells; this model assumes that the fraction of RNA-seq reads in the tumor sample that arise from cancer cells is proportional to tumor purity, and that the ASE of genes within normal cells within the tumor is the same as their ASE in the paired normal cells. In the third model, diffASE-exp, we adjust our differential ASE estimates to account for differential expression between cancer and normal cells; this model assumes that normal cells in the tumor sample have the same expression level for the gene on average as in the paired normal. For each model, we compute a per-site *p*-value to determine if the computed fraction of reads with the specific nucleotide in cancer cells is significantly different from the fraction in normal cells, given the number of reads estimated to arise from cancer cells.

**Figure 1:**
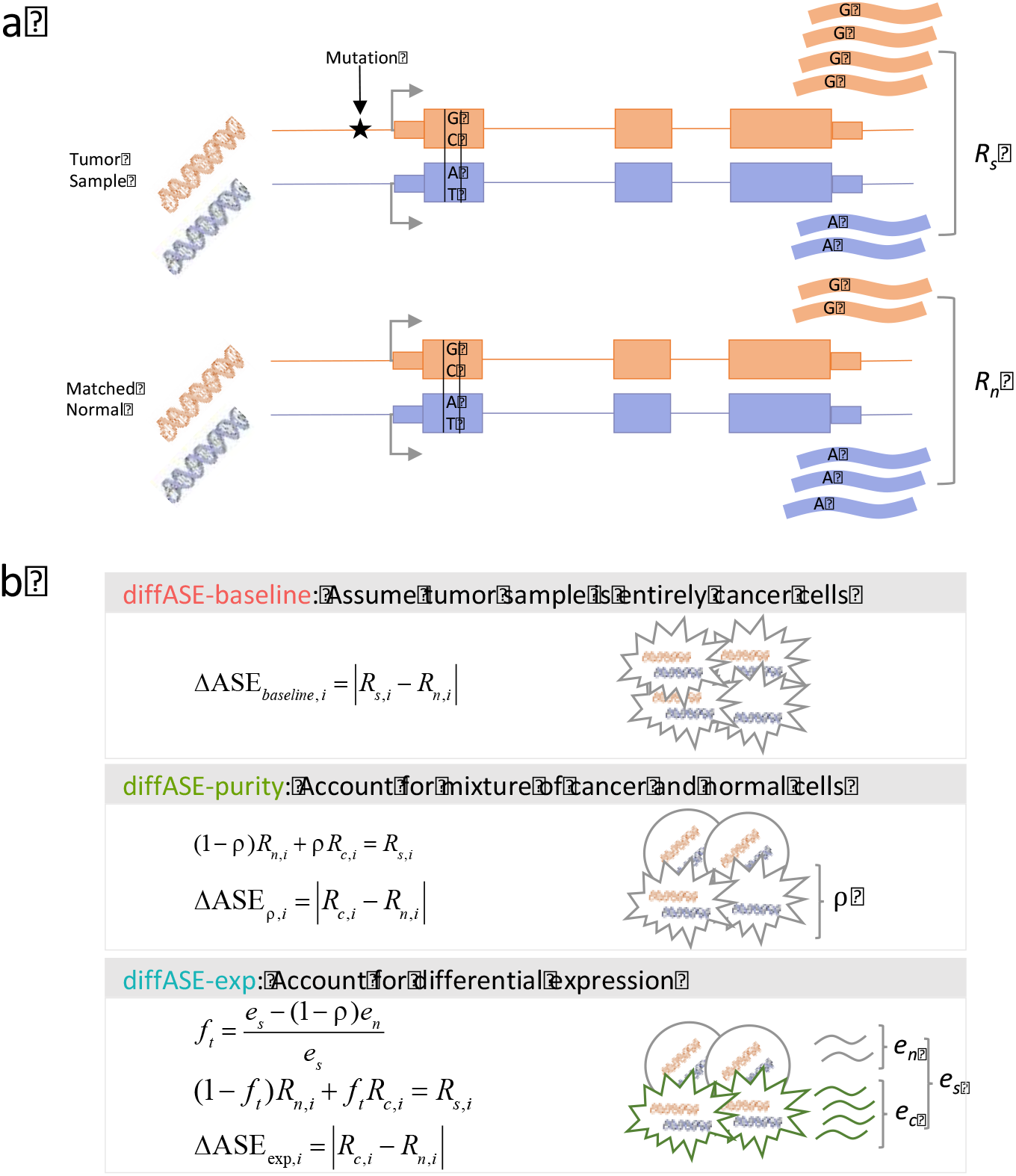
Overview of Differential Allele-Specific Expression Framework. **a.** The differential ASE of a transcript in an individual’s tumor considers both *R_s_*, the fraction of gene transcripts in the tumor sample that contain a particular allele, along with *R_n_*, the fraction in the matched normal sample. In this schematic, both the tumor and matched normal samples exhibit allelic imbalance. **b.** We develop three models to compute differential ASE at a heterozygous site *i*. Let *R_s,i_* and *R_n,i_* denote the fraction of all RNA-seq reads mapped to site *i* that have a given nucleotide at site *i* in the tumor sample and normal sample, respectively. The diffASE-baseline model (top) assumes that the tumor sample is entirely composed of cancer cells, and thus Δ*ASE_baseline,i_* is directly computed as the absolute difference between *R_s,i_* and *R_n,i_*. The diffASE-purity model (middle) accounts for tumor samples being a mix of cancer and normal cells by using a tumor purity estimate *ρ* to deconvolve RNA reads in the tumor sample into reads arising from cancer cells and normal cells. Assuming that ASE estimated from the normal sample is a good approximation for ASE in the normal cells within the tumor sample, we can solve for *R_c,i_*, the fraction of reads from cancer cells that contain the given nucleotide. Δ*ASE_ρ,i_* is then the absolute difference between *R_c,i_* and *R_n,i_*. The diffASE-exp model (bottom) extends this concept by accounting for differential expression between cancer and normal cells. We assume that the normal cells in the tumor sample are on average expressed at the same levels as in the matched normal. This allows us to compute *f_t_*, the fraction of RNA reads that are derived from cancer cells by subtracting away the ones that are derived from the normal cells in the sample. We then proceed as in the prior model using that fraction in the place of purity. For more details, see Methods.

Simulations (see Supplement, Section A) show that the models can successfully detect differential ASE in realistic settings for sample purity and read depth and that more complex models tend to better estimate differential ASE (Figures S1-S3). Simulations predict that differential ASE is easier to detect in cases where genes are overexpressed in cancer cells (Figure S2). Conversely, simulations indicate that, if read depth is low, differential ASE may be hard to detect when genes are underexpressed in cancer cells (Figure S3). We observe that our models are robust to variation in tumor purity (Figure S4) and to sequencing errors (Figure S5). As expected, when ASE is high in both normal and cancer cells, ASE estimates using just tumor samples (i.e., one-sample or tumor sample ASE) are poor estimates of the amount of differential ASE in cancer cells as compared to normal cells (Figure S6). Precision-recall curves computed on simulated data reveal that our models outperform tumor sample ASE and two-sample MBASED (Figures S7 and S8). In particular, our models have higher precision across almost all levels of recall, indicating that the adjustments for heterogeneity that our models introduce are critical to accurately identify differential ASE.

Because TCGA data is unphased, to obtain gene-level estimates of differential ASE using any of the three models, for each gene, we take a read depth weighted median value over all its heterozygous sites. Per-gene significance values are obtained by combining *p*-values across all sites within a gene using Fisher’s method and computing false discovery rates (FDRs) via the Benjamini-Yekutieli procedure (Benjamini & Yekutieli, 2001). See Methods for more details.

### The Landscape of Differential Allele-Specific Expression in Breast Cancer

In the TCGA breast cancer data set, 91 individuals have matched RNA-seq data for both tumor and normal samples, along with copy number variation (CNV) and whole-exome sequencing data. Since immune cells can infiltrate tumors and may confound our analyses (as we assume that non-cancer cells in a tumor sample are modeled well by paired normal tissue samples and can be deconvolved based upon estimates of tumor purity and expression), we used Cibersort (Newman et al., 2015) to remove tumor samples with significant expression correlation to immune cell types (*p*-value < 0.05). We ran the models on all genes for the remaining 46 individuals as described in Methods. While there are many gene-sample pairs that are considered to exhibit significant differential ASE by all three of our models, each model selected a slightly different set as significant across the individuals (Figure 2a). Though simulations suggest diffASE-purity and diffASE-exp are more accurate than diffASE-baseline (Figures S1-S3), when analyzing each set of genes found to be significant by only one of our three models, we find that the three methods have complementary strengths. The diffASE-baseline model is able to detect differential ASE at lower read depths as compared to diffASE-purity; that is, among genes found to be significant by only diffASE-baseline, the median read depth per heterozygous site within these genes is 42 whereas this number is 57 for diffASE-purity. We also observe that diffASE-exp is better able to detect differential ASE in genes that are underexpressed in tumor as compared to normal (median log_2_-fold change in expression for genes found to be significant is −0.4 for diffASE-exp versus 0.5 for those found significant by either of the other two models). We thus chose to incorporate results from all three models to achieve the highest coverage of genes with significant differential ASE.

**Figure 2:**
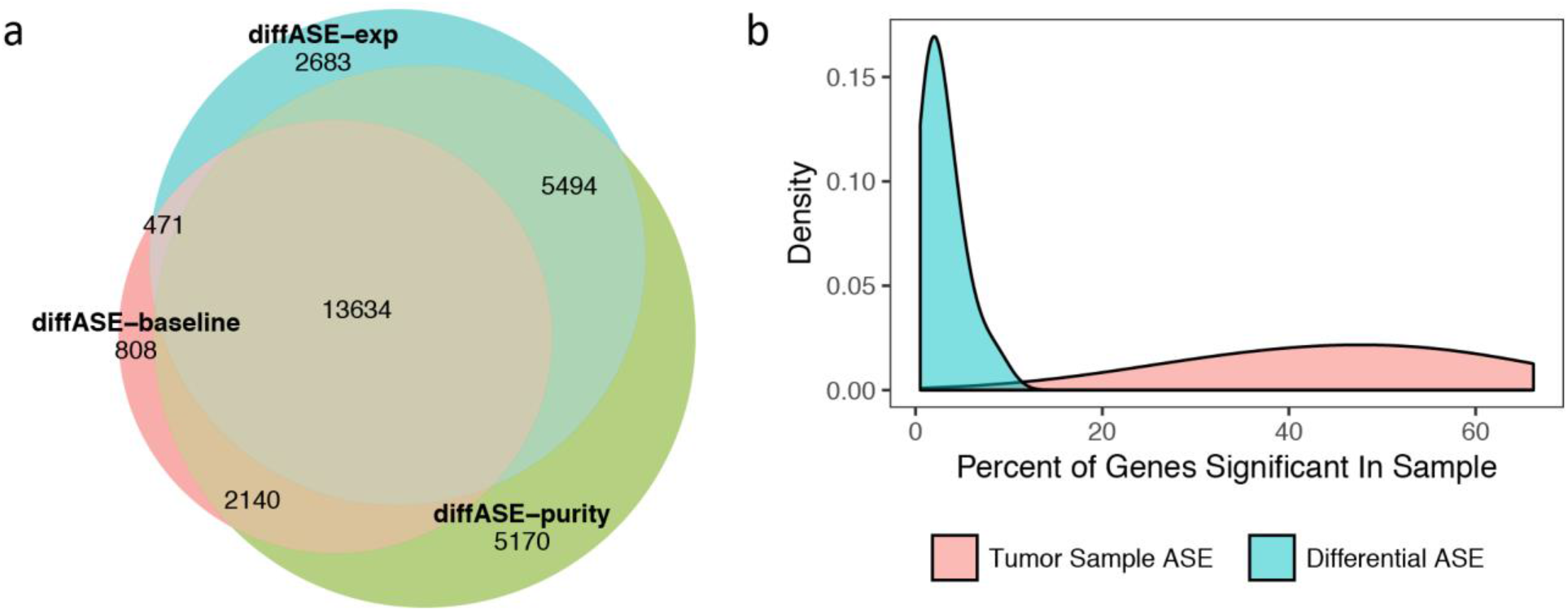
Landscape of Differential Allele Specific Expression in Breast Cancer. **a.** Numbers of gene-sample pairs with significant differential ASE using each of the three methods. While there are many gene-sample pairs that are called significant by all three models, each model selected a slightly different set of genes as significant across the individuals. **b.** The distributions of genes exhibiting significant differential ASE per individual when estimating differential ASE between cancer and normal cells using any of the three models (blue) versus tumor sample ASE computed using tumor sample RNA-seq data only (red). The median fraction of genes with at least one heterozygous site that exhibit significant ASE is 47.0%, whereas the median fraction of these genes exhibiting differential ASE is only 2.18%, indicating that the majority of ASE observed in a tumor sample by the conventional method is likely also present in the matched normal. For both panels, in order to be called significant, a minimum of differential ASE (or ASE) of > 0.15 is required along with an FDR < 0.1.

We consider a gene to exhibit significant differential ASE in an individual if in at least one model it has an FDR < 0.1 and an estimated differential ASE > 0.15. We found that across individuals, the median fraction of genes where ASE could be determined (i.e., with at least one heterozygous site) that exhibit significant differential ASE is 2.18%. In contrast, across individuals, a median of 47.0% of these genes are significant when computing ASE on tumor samples alone with the same FDR and ASE effect size thresholds (Figure 2b). This is consistent with previous studies (Ma et al., 2018; Mayba et al., 2014; Romanel et al., 2015), as we can only compute significance in, on average, 11,585 genes per individual as the remaining genes do not contain at least one heterozygous site. The discrepancy between the number of significant genes when using tumor sample ASE versus differential ASE indicates that the majority of ASE observed in a tumor sample by the conventional method is likely also present in the matched normal. In contrast to differential expression analysis, which considers a collection of samples and typically uncovers numerous differentially expressed genes between tumor and normal samples (Koboldt et al., 2012), differential ASE has specificity in detecting regulatory events in cancer and is able to detect per-individual regulatory events.

While a gene may exhibit differential ASE due to somatic alterations in its regulatory regions, it may also arise for other reasons, including nonsense-mediated decay, aberrant methylation, or CNVs where amplifications or deletions of the gene on one copy lead to changes in expression for that copy. These situations provide a useful validation for our approach to uncover differential ASE. CNVs in particular have a large impact on ASE (Mayba et al., 2014). We observe that genes affected by CNVs have higher differential ASE than other genes (Supplement, Section B) and are enriched in the set of gene-sample pairs with significant differential ASE (one-sided hypergeometric *p*-value = 6.12e-37), confirming that our models can detect expected allele-specific changes in expression resulting from amplifications and deletions. Further, we find that, when considering genes with at least one heterozygous site in a sample, a per-sample median of 4.31% of genes with CNVs are significant while only 1.83% without CNVs are significant. Our approach appears to be equally effective at detecting differential ASE arising from either amplifications or deletions: across samples, of genes with CNVs, a median of 4.46% of genes within regions of amplification and 4.22% of genes in regions containing deletions are significant (Figure 3a). While differential ASE is harder to uncover for underexpressed genes (Figure S3), this is mitigated by the large impact of CNVs on ASE (Figure S5).

**Figure 3:**
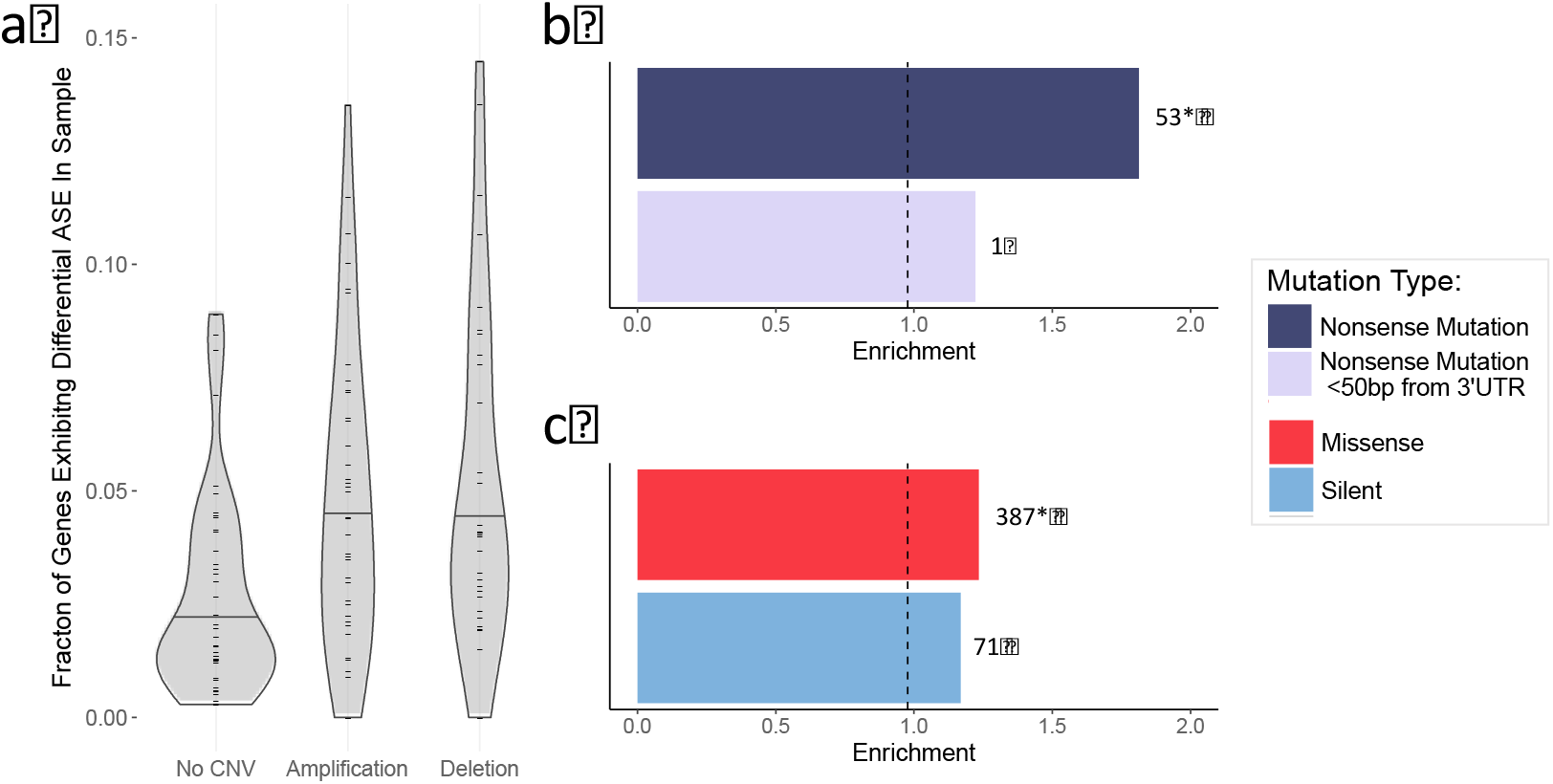
Differential ASE Detects Known Causes of ASE. **a.** The fractions of significant genes per individual when the genes contain amplifications, deletions, or no copy number variants. Across individuals, a median of 4.46% and 4.22% of genes with CNV amplifications and deletions, respectively, are significant while only 1.83% without CNVs are significant, confirming that our model can detect the expected allele-specific change in expression resulting from CNVs. **b.** Enrichment of gene-sample pairs that have nonsense mutations (dark purple) and the subset of nonsense mutations within 50bp of the 3’UTR (light purple) amongst all gene-sample pairs that exhibit significant differential ASE. For each mutation type, to the right of the bar, the number of gene-sample pairs that exhibit differential ASE and have that type of mutation is given. Enrichments with hypergeometric *p*-values < 0.05 are starred. Whereas there is a significant 1.81-fold enrichment for all nonsense mutations, nonsense mutations within 50bp of the 3’UTR exhibit a non-statistically significant 1.22-fold enrichment. Gene-sample pairs with no mutations have no enrichment amongst all genes with significant differential ASE (dashed vertical lines at 0.98). **c.** Gene-sample pairs with significant differential ASE are also enriched for missense mutations (red) and silent mutations (blue), but this enrichment is only significant for missense mutations.

### Differential Allele-Specific Expression Captures Effects of Coding Mutations

To examine the impact of mutations on differential ASE, we looked at coding mutations occurring in gene-sample pairs unaffected by CNVs. Nonsense mutations are expected to reduce the number of transcripts from the copy of DNA they appear on due to nonsense-mediated decay (Lindeboom et. al, 2016; Rhee et. al, 2017). We find that genes-sample pairs that exhibit significant differential ASE are 1.81-fold enriched for nonsense mutations as compared to the baseline fraction of all gene-sample pairs containing nonsense mutations (Figure 3b, dark purple), which represents a statistically significant level of enrichment (one-sided hypergeometric *p*-value = 6.67e-5). As a comparison, significant gene-sample pairs are 0.98-fold under-enriched for those pairs that do not have mutations in exome sequencing data (Figure 3b, dashed line). Furthermore, it has been shown that nonsense-mediated decay is less likely to occur if a nonsense mutation occurs within 50 bases of the 3’UTR (Lindeboom et al., 2016; Rhee et al., 2017). In agreement with this, of the 28 gene-sample pairs where such mutations are observed, we find only one with significant differential ASE; this is a 1.22-fold enrichment for such mutations (Figure 3b, light purple) and is not significantly different from random (one-sided hypergeometric *p*-value = 0.17).

Next, we considered the effect of missense and silent mutations on differential ASE. The effect of these two mutation types in cancer on ASE has previously been observed (Diederichs et al., 2016; Rhee et al., 2017). In order to examine the effect of each given mutation type exclusively, we only consider gene-sample pairs that contain that type of mutation in an evolutionarily conserved site and no other mutations in conserved sites, except for intron mutations, of which there are too many of to exclude. We find that there is a modest 1.23-fold enrichment for gene-sample pairs with missense mutations (Figure 3c, red) and a lower 1.17-fold enrichment for gene-sample pairs with silent mutations (Figure 3c, blue) among all gene-sample pairs exhibiting significant differential ASE. The enrichment for gene-sample pairs with missense mutations is significant (one-sided hypergeometric *p*-value = 3.29e-4) while the enrichment for gene-sample pairs with silent mutations is not (one-sided hypergeometric *p*-values = 0.064). Interestingly, among all gene-sample pairs with significant differential ASE, the fraction with missense mutations does not differ significantly from the fraction with silent mutations (one-sided Binomial *p*-value = 0.39), suggesting that these mutations may affect ASE by impacting transcript stability.

### Known Cancer Genes Recurrently Exhibit Differential Allele-Specific Expression

One of the most powerful techniques to identify cancer-relevant events is to consider recurrence across samples so we next examined genes that recurrently exhibit significant differential ASE across samples. A gene recurrently exhibiting differential ASE at least six times across samples is statistically significant (Benjamini-Hochberg corrected one-sided Poisson binomial FDR < 0.1). Due to the strong effect of CNVs, we examined recurrence separately across genes without CNVs and those with, in both cases finding many cancer-related genes (Figure 4a, genes ordered by non-CNV recurrence on the left and by CNV recurrence on the right). Among those without CNVs, among the top 10 most recurrent genes, two are known cancer genes from the Cancer Gene Census (Futreal et al., 2004), a significant enrichment (hypergeometric *p*-value = 0.02), and seven show significant differential expression between cancer and normal when considering only the set of individuals that exhibit significant differential ASE in that gene, despite the fact that significance is difficult to obtain with such small numbers of samples. We find that the oncogene *MDM2* is significantly overexpressed in samples showing significant differential ASE (edgeR (Robinson et al., 2009) *p*-value = 1.5e-3), indicating that the observed dysregulation may be contributing to cancer development. Among the other genes, *RABEP1* promotes breast cancer growth (Yan, Xu, Tan, Liu, & He, 2011), *GSR* is involved in oxidative stress response, *RPA1* is a component of DNA repair (O’Leary et al., 2016), *MREG* is a functional partner of the *RAB27B* oncogene (Szklarczyk et al., 2017), and *NUP88* has been linked to cancer progression (Agudo et al., 2004). Among the recurrently significant genes with CNVs, *NAT1* is known to activate carcinogens, and all six recurrences are amplifications (Szklarczyk et al., 2017), while *ASAH1* is thought to play a role in cancer progression, and all six recurrences are deletions (O’Leary et al., 2016). *ASAH1* additionally exhibits significant differential ASE in four samples that do not have CNVs in that gene. *KMO* is thought to play a role in cancer by preventing apoptosis, and all six recurrences are amplifications (Wilson et al., 2016). *NSL1* is an important component of cell division and interacts with *CASC5* (Cancer Susceptibility Candidate 5) (Szklarczyk et al., 2017). Finally, *PRUNE* is known to play a role in cell proliferation, and all six recurrences are amplifications (O’Leary et al., 2016). It also exhibits significant differential ASE in two additional samples without CNVs.

**Figure 4:**
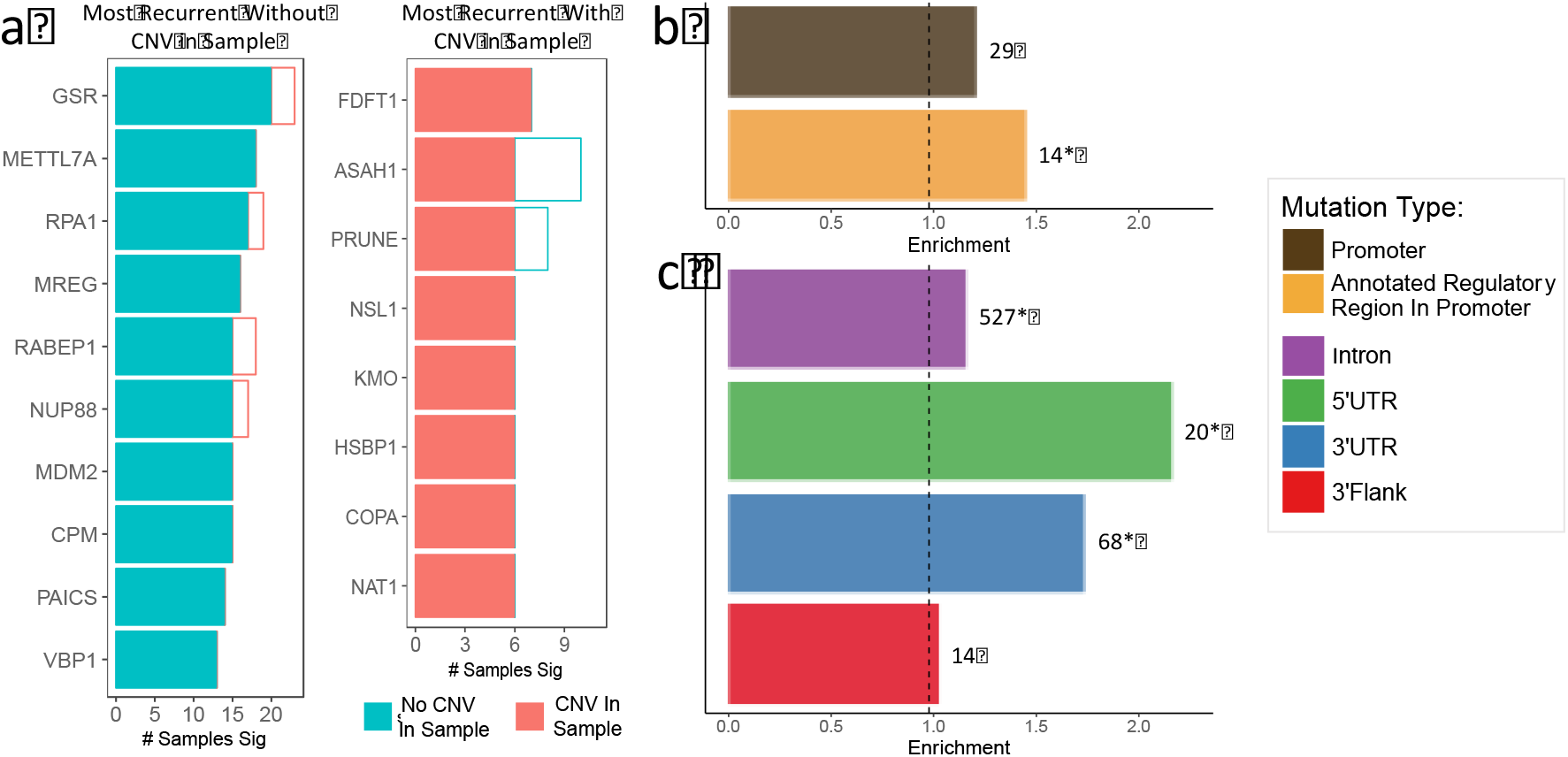
Regulatory Changes are Detected by Differential ASE. **a.** Genes that most frequently exhibit differential ASE when ordered by recurrence across samples without CNVs (left) and ordered by recurrence across samples with CNVs (right). In both cases, the count for samples without CNVs is in blue and the count for samples with CNVs is in red. **b.** The enrichment (see Figure 3 caption) among gene-sample pairs exhibiting significant differential ASE for promoter mutations (brown) and the subset of promoter mutations that are annotated to regulatory regions (orange) as compared to no mutation (dashed line). **c.** The enrichment among gene-sample pairs exhibiting significant differential ASE for mutations in an intron (purple), 5’UTR (green), 3’ flank (red), or 3’ UTR (blue) as compared to no mutation (dashed line).

To evaluate the list of all genes recurrently exhibiting differential ASE (for the full list, see Supplement, Section C), we looked for enrichment in cancer-related gene sets. We used gene set enrichment analysis on the list of genes ranked by recurrence of differential ASE across samples (not counting samples with CNVs in those genes, as we wish to exclude the functional effects of CNVs). We considered all pathways from Reactome as well as all diseases from the Disease Ontology (Yu & He, 2016). The most significant pathway is “stabilization of p53,” an important pathway for cancer development. Other relevant significant pathways include “apoptosis” as well as specific cancer-related pathways such as “Autodegradation of Cdh1” and “APC-Cdc20 mediated degradation of Nek2A” (for a full list of significant pathways see Supplement, Section D). The top nine Disease Ontology gene sets are all cancers, and breast cancer appears in the full list of significant terms (for a full list of significant disease ontologies, see Supplement, Section D). These results indicate that genes recurrently exhibiting differential ASE are broadly involved in cancer processes.

Since breast cancer is subtyped partly based on the overexpression of estrogen receptors, we examined differential ASE in subtyped samples to determine the cause of dysregulation in those samples, as noncoding mutations may play a role (Bailey et al., 2016). Among the 22 individuals who were clinically subtyped as estrogen receptor positive (ER+) and had higher levels of alpha estrogen receptor gene (*ESR1*) in their cancer cells than in their normal cells, five individuals exhibited significant differential ASE for ESR1, while another two had amplifications due to a copy number alteration; this suggests that for a notable fraction of ER+ individuals, the overexpression of their *ESR1* gene is due to cis-regulatory alterations. Further, all samples with significant differential ASE in *ESR1* overexpressed it. We note that amongst the ER-patients, none exhibited differential ASE for ESR1. Finally, to determine if dysregulation was broadly similar among ER+ patients, we computed the distances between all individuals across genes by measuring the Euclidean distance between a binary vector for each individual of whether each gene was significant with respect to differential ASE or not in that individual. We find that the differential ASEs across genes between ER+ individuals are significantly more similar to each other than to ER-individuals (Wilcoxon *p*-value = 6.56e-6) suggesting that similar sets of genes are altered in *cis* across individuals within subtypes.

### Functional Noncoding Mutations Lead to Differential Allele-Specific Expression

To measure the impact of noncoding mutations on differential ASE, we looked for enrichment in the presence of promoter, 5’UTR, 3’UTR, 5’Flank, and intronic mutations among gene-sample pairs exhibiting significant differential ASE. As with coding mutations, in order to examine the effect of each given mutation type exclusively, we only consider gene-sample pairs that contain that type of mutation in an evolutionarily conserved site exclusively, with the exception of mutations in introns. We find that gene-sample pairs with promoter mutations are 1.23-fold enriched in gene-sample pairs exhibiting significant differential ASE (Figure 4b, brown), which is not a statistically significant enrichment given the number of such mutations (one-sided hypergeometric *p*-value = 0.079). However, if we filter the set of promoter mutations to only those falling in sites annotated as regulatory by the curated site ORegAnno (Lesurf et al., 2016), we observe a much larger 1.44-fold enrichment (Figure 4b, orange) that is statistically significant (one-sided hypergeometric *p*-value= 0.034). Moreover, when we restrict to promoter mutations that fall into sites that are not annotated as regulatory, we see only 1.06 fold enrichment (hypergeometric *p*-value= 0.25). When we consider gene-sample pairs with a mutation in their 5’UTR, we observe the strongest enrichments within gene-sample pairs with significant differential ASE (2.16-fold enrichment, one-sided hypergeometric *p*-value = 4.59e-4, Figure 4c, green). We also observe a large enrichment for differential ASE in gene-sample pairs with mutations in the 3’UTR (1.73-fold enrichment, one-sided hypergeometric *p*-value = 1.32e-5, Figure 4b, blue), a small but significant enrichment for gene-sample pairs with mutations in their introns (1.16-fold enrichment, one-sided hypergeometric *p*-value = 2.56e-4, Figure 4c, purple), and no enrichment for those with mutations in the 3’Flank (1.03-fold enrichment, one-sided hypergeometric *p*-value = 0.20, Figure 4c, red). In contrast, when using tumor sample ASE, of the noncoding mutations, only 3’UTR mutations had a very small but significant enrichment among gene-sample pairs that are significant (Supplement, Section E), indicating that differential ASE is crucial to measuring the effects of noncoding mutations.

One of the most powerful features of differential ASE is that it serves as a means to distinguish which *cis* noncoding mutations have the potential to cause dysregulation. Mutations *cis* to genes that have no observed differential ASE are most likely not leading to dysregulation of that gene and thus can be filtered out as potential functional noncoding mutations. This is important because there are numerous noncoding mutations in any given sample, most of which have no functional impact. Among the samples we studied, there are 23,400 noncoding mutations observed in whole-exome sequencing data in conserved sites. However, only 769 of these are *cis* to genes with observed differential ASE (for a full table of counts by mutation type see Supplement, Section F). Filtering out 96.2% of mutations represents an enormous decrease in the search space for functional noncoding mutations. In comparison, tumor sample ASE is only able to filter 46.3% of conserved noncoding mutations.

To generate a list candidate functional noncoding mutations, we consider only those that occur *cis* to genes with significant differential ASE and occur within a regulatory site as annotated by ORegAnno (Lesurf et al., 2016). We note that multiple mutations can be implicated by the same gene, and thus to minimize alternative causes of dysregulation, we exclude any mutation for which the sample had a CNV or nonsense mutation in the same gene. These stringent requirements give us a list of nine promoter mutations, 15 5’UTR mutations, 15 3’UTR mutations, three 3’Flank mutations, and 105 intronic mutations (for a full list of mutations see Supplement, section G). Among these noncoding mutations, 5.6% are associated with known cancer genes from the CGC (Futreal et al., 2004), which is significantly enriched relative to the background rate of CGC genes (hypergeometric *p*-value = .011). This list also includes many other cancer relevant genes such as *PGRMC1* (Neubauer et al., 2013), *RHOB* (Chen et al., 2016), *HYOU1* (Krȩtowski et al., 2013), *INPPL1* (O’Leary et al., 2016), *GADD45B* (O’Leary et al., 2016), and *CDKN1B* (Chu et al., 2007).

We next sought to determine if there were potential functional mechanisms linking the effect of the observed mutations to changes in ASE. We tested all our mutations for their predicted regulatory effect using deepSEA (Zhou & Troyanskaya, 2015). We find that 22% of promoter mutations, 47% of 5’UTR mutations, 27% of 3’UTR mutations, two of the three 3’Flank mutations, and 8.6% of intronic mutations have a significant predicted functional impact. This indicates that a large proportion of our candidate mutations have the potential to affect gene regulation. We additionally used FIMO (Grant et al., 2011) with a set of transcription factors from HOCOMOCO (Kulakovskiy et al., 2018) to test if our set of mutations could disrupt or create new transcription factor binding sites. We found that eight of the nine promoter mutations cause at least one change in transcription factor binding sites. These results indicate that a significant fraction of the mutations we observe *cis* to genes showing significant differential ASE have the potential to cause the observed dysregulation.

Finally, to assess the power of differential ASE as a method for whole-genome patient-specific analysis, we examined mutations in the promoters of four samples that have whole-genome sequencing data available. While these individuals are also included in the whole-exome sequencing data, using whole-gene sequencing allows us to look further upstream from genes to detect additional mutations. Among these four samples, we find 451 promoter mutations in conserved sites within 2000 bases upstream of transcription start sites. Only 10 of these mutations are upstream of genes with differential ASE. When we limit these mutations to known regulatory regions as annotated by ORegAnno, we observed five potentially functional mutations (for a full list of mutations see Supplement, Section H). These appear upstream of KIF1B, TNFAIP8, PNN, FCF1, and PUDP, the first three of which have previously been associated with cancer (Day et al., 2017; Vitiello et al., 2014; Yang et al., 2016). Furthermore, four out of these five mutations are predicted by deepSEA to have a significant functional impact. Thus, the powerful filtering provided by differential ASE makes it possible to focus on only the subset of mutations that are potentially functional in whole genome sequencing.

## Discussion

We have shown that differential ASE is a powerful tool for honing in on genes that are dysregulated due to somatic *cis* alterations. We have demonstrated that genes that recurrently exhibit differential ASE across samples are cancer related, and that we can detect the effects of CNVs, nonsense-mediated decay, coding mutations, and noncoding mutations. While ASE has previously been analyzed in cancer genomes, our work suggests that many genes exhibit ASE in cancer cells but that most of these do so due to their expression patterns in normal cells. One sample ASE by itself is thus not an effective means to analyze cancer genomes.

Differential ASE analysis identifies 147 potentially functional noncoding mutations without relying on recurrence across individuals. While these mutations are strong candidates in terms of functional impact, they do not overlap noncoding mutations that have been previously proposed as functional in breast cancer (Corces et al., 2018; Nik-Zainal et al., 2016; Rheinbay et al., 2017; Vinagre et al., 2013; Y. Zhang et al., 2018). However, an examination of *all* noncoding mutations observed in the exome sequencing data of the individuals included in our study reveals that only one of the previously identified candidate functional noncoding mutations occur in *any* of these data. Interestingly, we did observe significant differential ASE in the associated nearby gene *NFKBIZ* in the individual with that noncoding mutation, but the mutation itself did not pass our filters (i.e., falling into a conserved site within an annotated regulatory region). Overall, the low amount of overlap is not surprising due to the low levels of recurrence of even the most well studied noncoding somatic mutations. As a result, the overlap in predicted functional mutations by different methods is expected to be small, and further development of methods such as ours that are not based on recurrence is critical.

We introduced three models to detect differential ASE, each of which takes a different approach to handle the heterogeneity observed in tumors. These three models uncover complementary sets of gene-sample pairs exhibiting differential ASE, and in the future, it may be possible to develop even more sensitive methods to deconvolve mRNA expression. For instance, not all cancer cells carry any given mutation. Models could account for the Variant Allele Frequency (VAF) of any given mutation to adjust the fraction of mRNA that is expected to derive from cells with that mutation. Further, models could consider tumor mutation phylogenetic trees, as earlier occurring mutations are more likely to be those that drive the progression of cancer (El-Kebir et al., 2016).

One important caveat to our work is that there are many events that can cause expression dysregulation and we were only able to consider a small fraction of them. For example, genes that recurrently exhibit differential ASE may be subject to distal *cis* regulatory events; the use of known enhancers or Hi-C contact maps may be helpful in constraining the sets of mutations that need to be considered (Orlando et al., 2018). Alternatively, it is known that there are epigenetic factors that can lead to ASE (Ongen et al., 2014). Furthermore, even if we find that a gene is dysregulated and suggest observed *cis* mutations as potentially causative, there may be multiple detected *cis* mutations and additional analysis is necessary to detect which of these are causing the change in ASE (Y. Zhang et al., 2018). Alternatively, other mechanisms such as differential splicing between tumor and normal could also result in ASE. It is also possible that a *cis* mutation, while functional, may not affect ASE due to more complex regulatory circuits or other compensatory alterations, thereby leading to false negatives. Another complication is the possibility of allele-specific read mapping bias (Degner et al., 2009), but in the majority of cases we expect this effect to be compensated for by differential analysis. Despite these challenges, however, because of the large number of noncoding mutations in cancer genomes, it is vital to have techniques for filtering out those mutations that are unlikely to be playing a regulatory role; indeed, we have found that the vast majority of noncoding mutations near genes do not appear to alter their expression.

Differential ASE analysis can only be applied to genes within a sample that contain heterozygous sites. In the BRCA samples in TCGA, estimates of differential ASE can be made for 30 to 60% of genes. While the TCGA datasets are predominantly comprised of European individuals, in more genetically diverse populations, differential ASE can be estimated for a larger fraction of genes. Importantly, since differential ASE is computed for genes in a per-individual manner, the overall composition of the population analyzed does not affect our analysis.

Differential ASE analysis in cancer necessitates the use of a matched normal sample, and thus this analysis may be better suited to some cancer types than others. We chose to analyze TCGA breast cancer samples because they comprise the only large data set where, across the same individuals, there is matched tumor-normal RNA-seq data and copy number variant calling, along with either whole-exome or whole-genome sequencing. Just as sequencing the germline is essential for uncovering somatic alterations in cancers, further studies that obtain expression information for both tumor and matched normal samples would be especially helpful in order to facilitate methods such as ours that control for variation in the normal sample. Other large-scale cancer sequencing efforts such as ICGC have the potential to generate such data (International Cancer Genome Consortium, 2010). Looking further ahead, widespread analysis of differential ASE without matched tumor-normal samples may be enabled by single-cell sequencing, as it should be possible to identify normal cells within a tumor sample.

Overall, our results suggest that differential ASE is a powerful way to detect recurrent dysregulation and to hone in on function *cis* noncoding mutations. Our observation that the majority of ASE detected in tumor samples is also present in the matched normal indicates that differential analysis is fundamental to examining dysregulation in cancer. Finally, the detection of differential ASE does not rely on observing recurring mutations across samples, and thus we believe that it could be used as a powerful tool for personalized medicine by identifying patient-specific functional noncoding mutations.

## Methods

### Estimating per-site differential allele-specific expression

In each individual, for each gene, we wish to compute its differential ASE, or the difference between the gene’s ASE in cancer cells and its ASE in normal cells. Since TCGA data is unphased, we first estimate differential ASE separately for each heterozygous site within the gene, and then combine these estimates. We propose three models with increasing complexity for the task of estimating differential ASE at a particular heterozygous site *i* (for a fixed gene within an individual) using paired RNA-seq data from tumor and normal samples.

At each heterozygous site, we refer to the variant that appears more frequently in the normal sample transcripts as the primary nucleotide. Let *R_c,i_* be the fraction of all RNA-seq reads mapped to site *i* that contain the primary nucleotide in cancer cells in the tumor sample. We want to compare this to *R_n,i_*, the fraction of all RNA-seq reads from normal cells that contain the same nucleotide at site *i*. However, because the tumor sample is a heterogeneous mixture of normal and cancer cells, we can only directly compute *R_s,i_*, the fraction of all RNA-seq reads in the tumor sample contain that the primary nucleotide at site i. The three models we propose estimate *R_c,i_*, from *R_s,I_* in different ways and then use this value to compute differential ASE, or Δ*ASE*, as |*R_c,i_* − *R_n,i_*|.

For our baseline model, **diffASE-baseline**, we compute Δ*ASE_baseline,i_* by assuming that *R_c,i_* = *R_s,i_*.

Our next model, **diffASE-purity**, considers *ρ*, the purity of the tumor sample (i.e., the fraction of cells in a tumor sample that are cancer cells), and computes Δ*ASE_ρ,i_*. This model assumes that the fraction of RNA-seq reads at site *i* in the tumor sample that arise from cancer cells is proportional to *ρ*, and that the ASE of genes within normal cells within the tumor is the same as their ASE in the paired normal cells. That is,

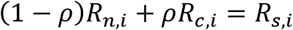

and we can solve for *R_c,i_* and then use it to compute Δ*ASE_ρ,i_*.

Our final model, **diffASE-exp**, additionally considers changes in expression for a gene between paired normal and tumor samples, and computes Δ*ASE_exp,i_*. This model uses tumor sample and normal expression counts for a gene (*e_s_* and *e_n_*, respectively, measured in counts per million) to estimate *f_t_*, the fraction of all transcripts for that gene in the tumor sample that arise from cancer cells. We assume that normal cells in the tumor sample have the same expression level for the gene on average as in the paired normal. We note that while it is likely that normal cells within the tumor sample have different expression from matched normal cells due to changes in their microenvironments, among other factors, they are expected to have more in common with each other than with tumor cells; that is, while imperfect, a gene’s expression within matched normal cells is a reasonable first approximation of its expression within normal cells in the tumor sample. With this assumption, the transcripts expressed within a tumor sample are a combination of normal transcripts and cancer transcripts adjusted for purity. That is,

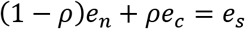

We can compute the fraction of transcripts for the gene in a tumor sample that arise from cancer cells as

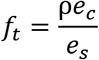

or equivalently that

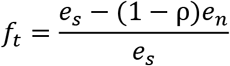

Then the RNA-seq reads in the tumor sample at site *i* within the gene arise from a mixture of normal and cancer transcripts weighted using *f_t_*

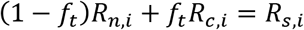

We can again solve for *R_c,i_* and use it to compute Δ*ASE_exp,i_*.

### Estimating the per-site significance of differential ASE

For a heterozygous site *i* within an individual, we determine whether its estimated *R_c,i_* is significantly different from *R_n,i_* as follows. First, for each model, we estimate the total number of RNA-seq reads at site *i* arising from cancer cells, *T_c,i_*, according to the assumptions of each model. For the diffASE-baseline model, we assume that *T_c,i_* is equal to the number of mapped reads at site *i* in the tumor sample. For the diffASE-purity model, we use *ρ* times the number of reads in the tumor sample. For the diffASE-exp model, we use *f_t_* times the number of reads in the tumor sample. Next, for each model, using the estimated *R_c,i_* and *T_c,i_*, we compute the number of reads with the primary nucleotide arising from cancer cells, and determine the probability of observing this number or a more extreme value. Our null hypothesis is that *R_c,i_* = *R_n,i_*, and we use the normal approximation of the binomial distribution with a mean of *R_n,i_T_c_* and a variance of *R_n,i_(1 − R_n,i_)T_c,i_*. In other words, the null hypothesis we are testing for each site is whether the number of reads of the primary nucleotide observed in cancer cells, estimated based on each model’s adjustments for purity and expression, is not different from the number that would be expected based on the primary nucleotide frequency in the matched normal sample.

### Per-gene estimates of differential ASE

For a given gene in an individual, for each model, we compute two gene-level measures, one for statistical significance and one for effect size. For each individual, we only consider genes with at least one heterozygous site. To compute statistical significance for each gene, we combine the *p*-values at each heterozygous site using Fisher’s method. Because sites with very high read depth would dominate this measure, if a site has more than 100 reads mapped to it in the tumor sample, we downsample to a maximum of 100 reads. We note that downsampling results in more conservative estimates of significance because fewer reads means a decrease in the power of the statistical test. If a gene has more than 10 heterozygous sites, we randomly sample ten of them ten times and take the median *p*-value of these randomizations, again generating a more conservative estimate of significance. To estimate a gene-level differential ASE value, we take a read depth weighted median of per site differential ASE, also considering a weight of at most 100 if there were more than 100 reads at a site.

For a particular method to estimate differential ASE, we say a gene has significant differential ASE if it is statistically significant at an FDR < 0.1 (Benjamini–Yekutieli method (Benjamini & Yekutieli, 2001)) and has a gene-level estimated differential ASE > 0.15. The median number of significant genes changes by fewer than ten genes per sample for FDRs between 0.05 and 0.2 and weighted differential ASEs between 0.1 and 0.2 (Supplement, Section I). Finally, to combine models, we consider a gene to be significant overall if is significant in at least one model. We note that we found no statistically significant difference between gene-sample pairs with significant differential ASE and those without with respect to of the number of heterozygous sites across the length of the gene (one-sided Wilcoxon *p*-value = 0.86).

### TCGA data acquisition and processing

In the TCGA (TCGA Research Network) breast invasive cancer (BRCA) cohort, 91 individuals have matched RNA-seq data for both tumor and normal samples, along with copy number variation and exome sequencing data. We downloaded somatic mutation calls identified by MuTect2 (Cibulskis et al., 2013) using the exome data. Across the 91 individuals we analyzed, MuTect2 identified 103,748 coding mutations and 303,415 noncoding mutations (for a full table of mutation counts by type, see Supplement, Section F). Of the samples we examined, four additionally have whole genome sequencing (WGS) data. We called mutations in the WGS data using MuTect2 with default parameters. For the WGS data, we considered promoter mutations to be any occurring within 2000bp upstream of a gene.

We used the *TxDb.Hsapiens.UCSC.hg38.knownGene* package in R (Maintainer, 2016) to download the locations of all annotated human genes excluding those on the Y chromosome. For each of these 22,732 genes, for each individual we considered all RNA-seq reads from the solid tissue normal BAM files that mapped uniquely to that gene (mapq quality code 255). For RNA-seq data, in cases where two vials were processed for a sample, we always used vial A. For each individual and gene pair, we used the *AllelicImbalance* package in R (Gådin et al., 2015) to find every site in the gene that had at least 20 mapped reads of which the more frequent nucleotide is at most 90 percent, filtering out those sites that had a somatic mutation observed in that individual. These sites comprise our collection of heterozygous sites for the gene in the individual. Next, for each individual and gene pair, for each heterozygous site identified in the gene in that individual, we counted both the total number of reads at each heterozygous site in both tumor and normal, as well as a count for the number of times the primary nucleotide appeared in normal and that same nucleotide appeared in the tumor.

For each individual and gene pair, we used its tumor and matched normal expression count data as measured in count files to compute the corresponding expression values in Counts Per Million (CPM) using *edgeR* (Robinson et al., 2010) with library sizes normalized by the Trimmed Mean of M-values (TMM). Masked copy number segment files from TCGA for both tumor and matched normal samples were obtained. For each gene-individual pair, we calculated the average segmented copy number score across the gene for both tumor and normal samples. For an individual, a gene is identified as falling into an amplified (respectively, deleted) region in the tumor if the log2-fold change in average segmented copy number score is > 0.5 (respectively, < −0.5). For each tumor sample, purity estimates were obtained from Aran et al. (Aran et al., 2015). In particular, we used the combined purity estimate (CPE) when available, and for the samples where the authors did not provide a combined estimate, we used the median of the available purity measures that comprise their combined estimate.

For each tumor sample, we used the Cibersort webserver to determine the correlation of gene expression CPM values with the LM22 set of immune expression signatures (Newman et al., 2015). We filtered out individuals whose tumors had a significant correlation with immune expression profiles at *p*-values < 0.05, leaving us with 46 individuals. In the Supplement, we consider results when filtering out larger numbers of individuals by considering tumor expression correlations with immune expression profiles at less stringent *p*-value thresholds (Supplement, Section I). We note that increasing the threshold to 0.1 only removes one additional individual.

### Functional and genomic analysis

The Cancer Gene Census (CGC) from COSMIC (Futreal et al., 2004) consists of 595 genes and comprises our list of known cancer genes. We compute Gene Set Enrichment using the package *ReactomePA* in R (Yu & He, 2016). We compute differential expression using *edgeR* (Robinson et al., 2010). For our breast cancer subtype analysis, clinical data is obtained from the Broad GDAC Firehose (Broad Institute TCGA Genome Data Analysis Center, 2016), and we classified any individuals with “estrogen receptor status” that was “positive” as ER+ and any with a status of “negative” as ER-.

We queried the UCSC genome browser (Karolchik, 2004) using *ucscTableQuery* via the *rtracklayer* package in R (Lawrence et al., 2009) to determine 20-way phyloP conservation score at the location of each somatic mutation (Pollard et al., 2010). Regulatory regions as annotated by ORegAnno 3.0 were obtained from the UCSC genome browser (Lesurf et al., 2016). The functional impact of each noncoding mutation was assessed by the deepSEA webserver (Zhou & Troyanskaya, 2015) using the specific location and the base change of the mutation. We considered a mutation to have a functional impact if it had a functional significance score < 0.01. To determine if noncoding somatic mutations disrupt transcription factor binding sites, we ran FIMO (Grant et al., 2011) with default parameters using a 40 base region around each mutation against consensus binding models from the HOmo sapiens COmprehensive MOdel COllection (HOCOMOCO) version 11 (Kulakovskiy et al., 2018). We considered a mutation to change transcription factor binding if only one of either the germline sequence or the sequence resulting from the somatic mutation had a *p*-value < 1e-4 for a given motif.

We consider enrichment with respect to eight different sets of mutations (nonsense, missense, silent, promoter, 3’UTR, 5’UTR, 5’Flank, and intron, as annotated by TCGA). For mutations other than nonsense mutations, we only consider mutations occurring within a location with a phyloP score of at least 0.1 and exclude mutations from the set if there is any other conserved non-intronic mutation in the same sample *cis* to or in the same gene because it would be unclear which mutation is contributing to the observed effect. Throughout, enrichment for a specific mutation type is computed as the fraction of gene-sample pairs exhibiting differential ASE that have the mutation as compared to the baseline fraction of gene-sample pairs with that type of mutation.

### Lead Contact and Materials Availability

Further information and requests for resources and reagents should be directed to and will be fulfilled by the Lead Contact, Mona Singh. This study did not generate new unique reagents.

## Acknowledgements

This work was supported in part by NIH grants (R01-CA208148 and R01-GM076275 to MS) and by the NSF Graduate Research Fellowship Program (DGE-1656466 to PFP).

## Author Contributions

PFP and MS designed the study. PFP performed the analysis. PFP and MS wrote manuscript. All authors read and approved the final manuscript.

