## Supplemental Figures and Tables for "Differential Allele-Specific Expression Uncovers Breast Cancer Genes Dysregulated By *Cis* Noncoding Mutations"

**Supplemental Methods, Figures, and Tables For "Differential Allele-Specific Expression Uncovers Breast Cancer Genes Dysregulated By *Cis* Noncoding Mutations"**

**Section A**: Differential ASE on simulated data.
**Section B:** Differential ASE due to CNVs.
**Section C:** Full list of genes with recurrent differential ASE.
**Section D:** Full lists of pathways and disease ontologies found significant by Gene Set Enrichment Analysis.
**Section E**: Enrichment for mutation types when using tumor sample ASE.
**Section F:** Mutation counts in TCGA data and how many mutations are filtered out by differential ASE.
**Section G:** Full list of potential functional noncoding mutations.
**Section H**: Potential functional promoter mutations uncovered using whole-genome sequencing data.
**Section I:** Robustness of our results to parameter settings.

**Section A: Differential ASE on Simulated Data**

In order to assess the accuracy of our three models and compare them to tumor sample ASE, we generated simulated RNA-seq read counts for a heterozygous site within a mix of cancer and normal cells comprising a tumor sample. We can then compare the actual amount of differential ASE in cancer cells with the values estimated by the models.

Given a tumor sample of $n$ cells and assuming that the true purity of the sample is $\rho$, each of the $n$ cells is labeled as a cancer cell with probability $\rho$ or as a normal cell otherwise. Next, assume we have the true expression levels $e_c$ and $e_n$ of the gene in cancer cells and normal cells, respectively. We compute the amount of mRNA generated by each type of cell (normal or cancer) by multiplying the number of cells of that type by the corresponding expression level. Then, given the true fraction $R_c$ of the major allele in cancer cells (i.e., the fraction of transcripts expressing that allele, which is equivalent to the fraction of times the primary nucleotide is observed in RNA-seq data), the mRNAs arising from cancer cells are each randomly determined to contain the major allele with probability $R_c$ and the minor allele otherwise. Similarly, given the true fraction $R_n$ of the major allele in normal cells, the mRNAs arising from normal cells are randomly assigned the major allele with this probability. Altogether this now gives us a large pool of mRNA that reflects the tumor sample. Finally, we downsample from this pool of mRNA based on a given read depth (i.e., the total number of reads at that site).

Given a collection of cells, mRNAs, and alleles generated as described above, we directly compute an observed purity $\hat{\rho}$ (as we know which cells in the tumor sample are cancer and normal) and the fraction of the major allele in the tumor sample $\hat{R}_S$. Furthermore, we compute the fraction of transcripts in the sample that arise from cancer cells $\hat{f}_t$ directly from the generated mRNA. Thus, for each simulation we can compute a predicted level of differential ASE for each model, and compare it to the true differential ASE using the known $R_c$ the simulation was based upon, and we can compute a $p$-value of per-site significance, as described in Methods.

Our first goal is to assess the performance of our three methods to estimate differential ASE across three primary types of variation: true difference in allele-specific expression between tumor and normal cells, the number of RNA reads at the heterozygous site (i.e., RNA-seq read depth), and the magnitude of expression change between tumor and normal. To test how the models react to changes in differential allele-specific expression, we use three different levels of differential ASE: no differential ASE ($R_c = R_n$), low differential ASE ($|R_c - R_n| = 0.25$), and high differential ASE ($|R_c - R_n| = 0.5$). We test at high read depth (100 reads) and low read depth (25 reads) and with cancer cell expression levels at three times the rate and one-third the rate of normal. In our initial testing, we fix true purity to a typical value ($\rho = 0.8$) (Aran, Sirota, & Butte, 2015). For each model and condition, we measure the L2 error (or L2 loss) between the known true differential ASE used to generate the data and the observed value (computed as the sum over all sites of the squared difference between the true ASE and the observed ASE) as well as the percentage of times simulations were significant at $p$-value < 0.05. In additional testing, we estimate the robustness of our methods to changes in purity as well as to noise in RNA-seq reads at heterozygous sites.

Second, to assess the performance of our models in comparison to tumor sample ASE as well as to results obtained when running two-sample MBASED (Mayba et al., 2014), we generate simulated positive and negative sites of differential ASE and compute full precision recall curves.

We assume an equal number of sites exhibiting either differential ASE (positive examples) or no differential ASE (negative examples). In order to generate positives and negative examples, we first sample values of $R_n$ from a normal distribution with a mean of 0.7 and a standard deviation of 0.1, which approximates what we observe in the matched normal TCGA data. For negatives, we set $R_c = R_n$. For positives, we select a delta value (either 0.1 or 0.2) and set $R_c = R_n$ + delta or $R_c = R_n$ – delta (each with probability one-half). We then generate reads as described above.

**Figure S1** shows simulated results for no differential expression with **a.** high read depth and **b.** low read depth at varying levels of differential ASE. **Figure S2** shows the same but where the gene is overexpressed in cancer cells, and **Figure S3** where the gene is underexpressed. We note that in these cases there are virtually no false positives. **Figure S4** shows the robustness of our methods to changes in purity. **Figure S5** shows the robustness of our methods to sequencing errors. **Figure S6** shows a comparison of differential ASE to tumor sample ASE. **Figure S7** shows precision-recall curves for differential ASE and tumor sample ASE. **Figure S8** shows precision-recall curves for differential ASE and two-sample MBASED.

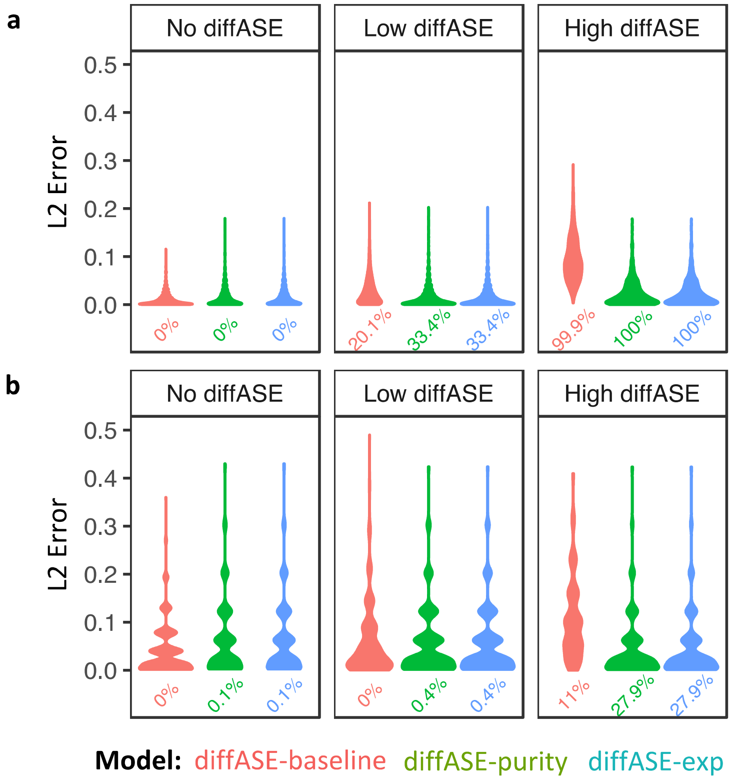


**Figure S1:** Results of simulations when there is no differential expression. Red represents diffASE-baseline, green diffASE-purity, and blue diffASE-exp. We ran 1,000 simulations in each setting. We show violin plots of the L2 error (y-axis) between the known true differential ASE used to generate the data and the value estimated by each model. The percentage under each distribution of L2 errors is the fraction of those runs found to be significant. **a.** Results with high read depth at varying levels of differential ASE. At high read depth, none of the models detect significance when there is no differential ASE (left) and always do at high differential ASE (right). With low differential ASE, we see that the more complex models have lower L2 errors and are more often able to detect significance. **b.** Results with low read depth at varying levels of differential ASE. At low read depth, the L2 error increases across the board as compared to high read depth. With high differential ASE, the more complex models have lower errors and are able to obtain significance nearly a third of the time. Overall, our simulations suggest that in the case where genes are expressed similarly in cancer and normal cells, high levels of differential ASE are readily and accurately detected at high read depths. Across the TCGA RNA-seq data we use, 80% of heterozygous sites have at least 20 reads, and 65% have at least 100, and therefore based on these simulation results, we are confident that differential ASE will be detectable in our data.

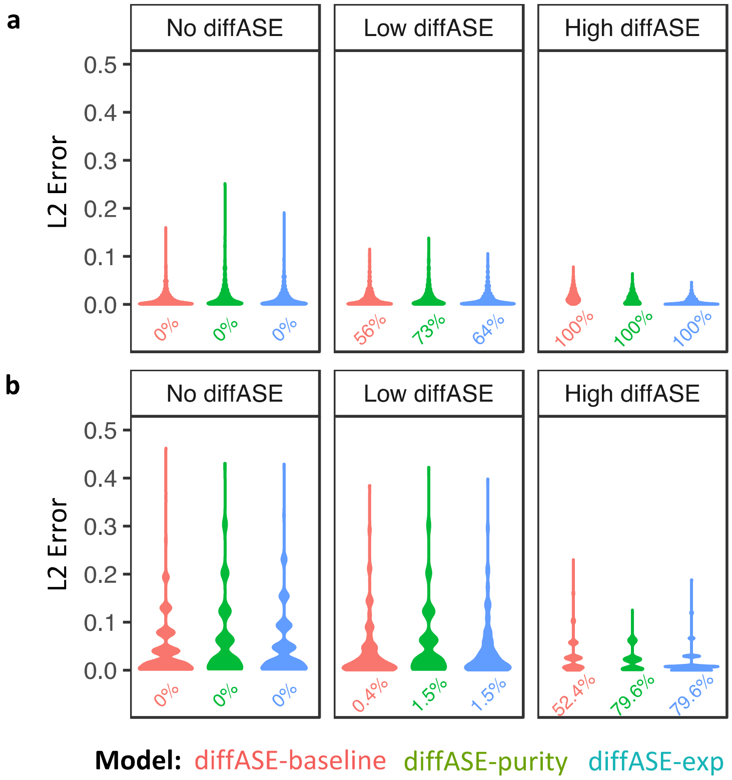


**Figure S2:** Results of simulations when a gene is overexpressed in cancer cells. As in Figure S1, red represents diffASE-baseline, green diffASE-purity, and blue diffASE-exp. We ran 1,000 simulations in each setting. We show violin plots of the L2 error (y-axis) between the known true differential ASE used to generate the data and the value estimated by each model. The percentage under each distribution of errors is the fraction of those runs found to be significant. **a.** Results for overexpression in cancer cells with high read depth and **b.** with low read depth at varying levels of differential ASE. Overall results are very similar to no differential expression (Figure S1 a). Error rates are lower and significance is easier to obtain across the board due to more reads being derived from cancer cells. Overall, our simulations suggest that when genes are overexpressed in cancer cells as compared to normal cells, differential ASE can be readily and accurately detected.

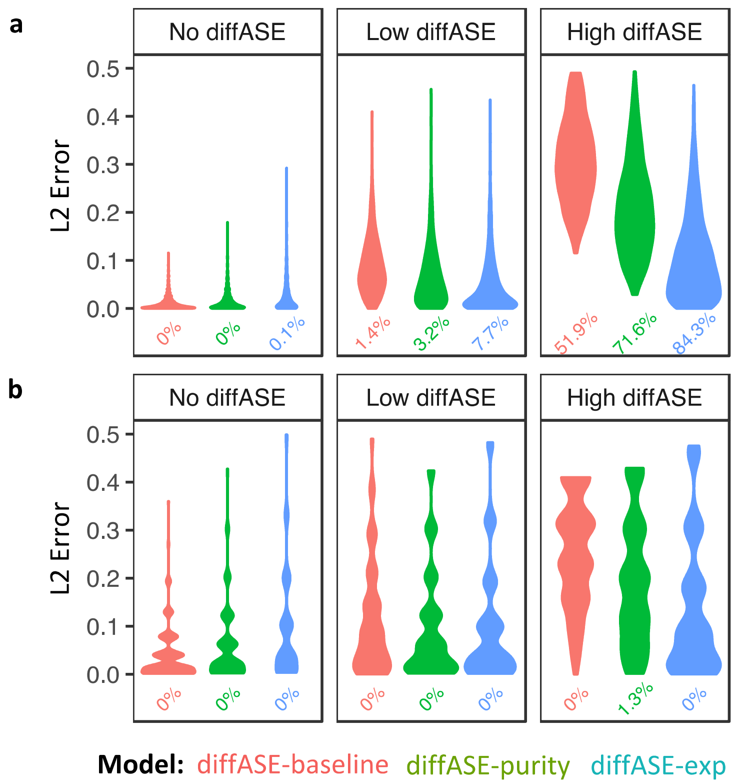


**Figure S3:** Results of simulations when genes are underexpressed in cancer cells. As in Figure S1, red represents diffASE-baseline, green diffASE-purity, and blue diffASE-exp. We ran 1,000 simulations in each setting. We show violin plots of the L2 error (y-axis) between the known true differential ASE used to generate the data and the value estimated by each model. The percentage under each distribution of errors is the fraction of those runs found to be significant. **a.** Results for underexpression in cancer cells with high read depth and **b.** with low read depth at varying levels of differential ASE. In the case when there is underexpression in cancer cells, fewer of the reads are derived from cancer cells and error rates increase as compared to the case where there are no changes in expression or when overexpression occurs in cancer cells. With high differential ASE with high read depth, more complex models can obtain significance at higher rates. However, at low read depth none of the models are able to obtain significance most of the times. Overall, our simulations suggest that when a gene is underexpressed in cancer cells as compared to normal cells, high read depth is needed, and only large amounts of differential ASE can be detected.

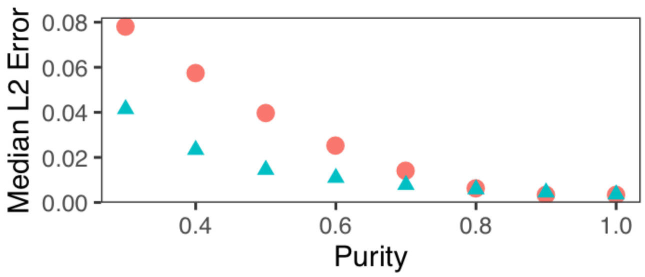


**Figure S4:** Robustness of our methods to changes in purity**.** We ran 1000 simulations at eight levels of purity between 0.3 and 1.0 (x-axis) at high read depth with $R_n$ = 0.5, $R_c$ = 0.7, and no differential expression. For purities between 0.8 and 1, diffASE-baseline (red circles) and diffASE-purity (blue triangles) have similarly small median L2 error between the known true differential ASE used to generate the data and the value estimated by each model. However, for purity lower than 0.8, diffASE-purity consistently has smaller errors than diffASE-baseline. As expected, as purity decreases, the error rates for both methods increase (e.g., at purity = 0.3, diffASE-baseline and diffASE-purity have median L2 errors of 0.04 and 0.08, respectively).

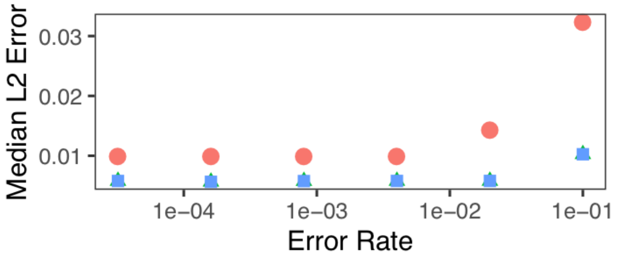


**Figure S5:** Robustness of our methods to sequencing errors. We introduce noise to our simulations by randomly changing between minor and major alleles at a heterozygous site with probability given by the error rate. We ran 1000 simulations at six error rates (x-axis, log scale) at high read depth and purity = 0.8, with $R_n$ = 0.5, $R_c$ = 0.7, and no differential expression. We find that for both diffASE-baseline (red circles) and diffASE-purity (blue squares), median L2 error between the known true differential ASE used to generate the data and the value estimated by each model does not begin to increase until the error rates is as high as 1 in 10, which is much higher than typically observed in such data. Thus, our methods are robust to errors in sequencing.

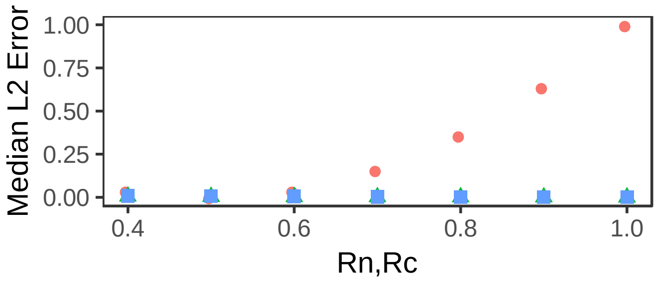


**Figure S6:** Differential ASE versus tumor sample ASE when the level of ASE is the same in cancer as in normal. The x-axis shows the level of ASE in cancer and normal, with the two values set equal to each other, while the y-axis shows the median L2 error between the known true differential ASE used to generate the data and the value estimated using tumor sample ASE (red circles), and diffASE-purity (blue squares). Results are across 1000 simulations at high read depth, purity = 0.8, and no differential expression. We do not show values estimated by diffASE-exp as that method is identical to diffASE-purity when there is no differential expression. We see that if the ratio in normal is the same as the ratio in cancer, as that fixed ratio changes, the error in tumor sample ASE estimates increases as the value moves away from 0.5. This indicates that, as expected, tumor sample ASE is unable to account for the level of ASE in normal cells.

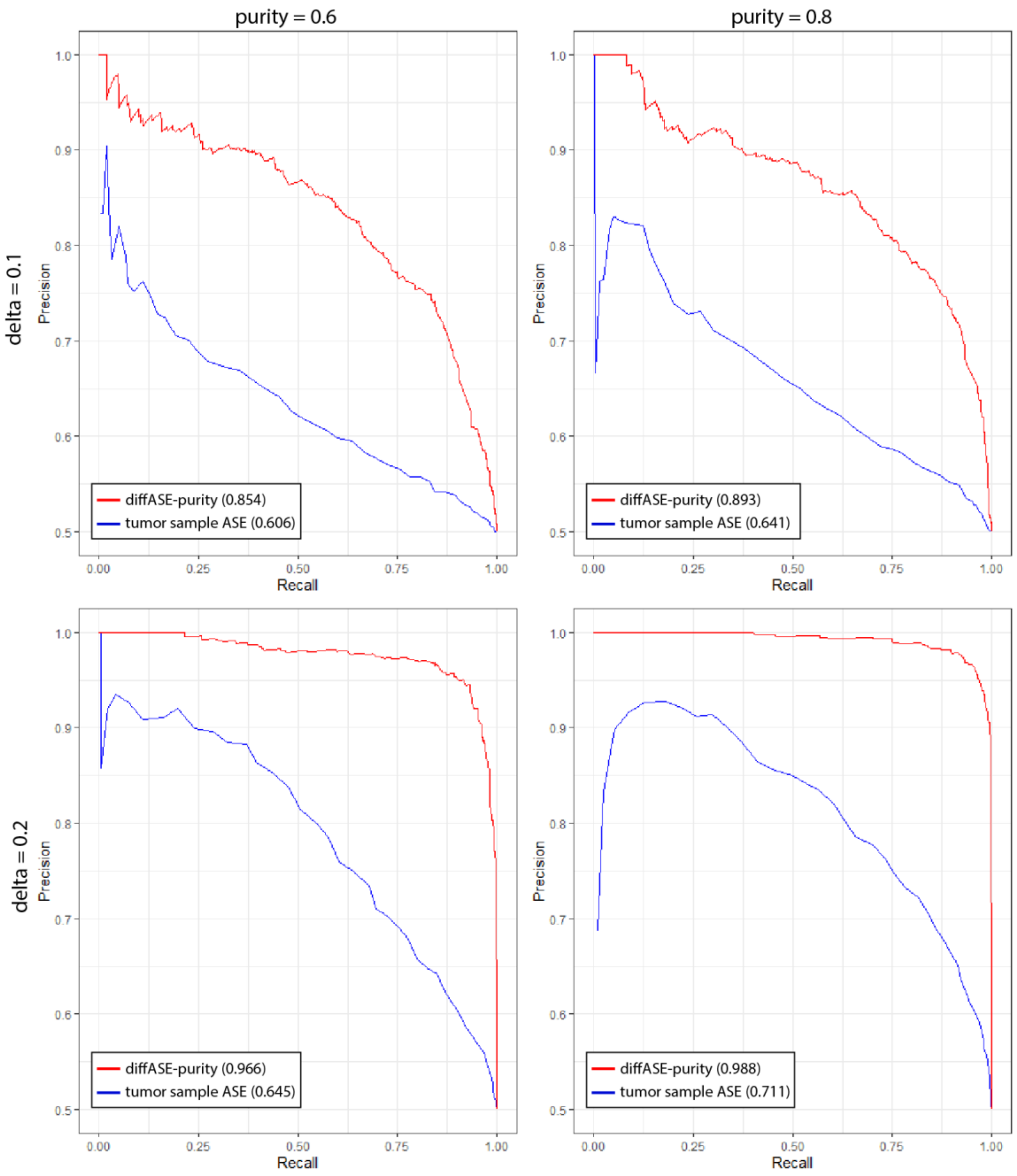


**Figure S7:** Precision-recall curves for differential ASE as determined by diffASE-purity and tumor sample ASE. We simulated 1000 negative and 1000 positive sites of differential ASE as described in Section A, with delta = 0.1 (top) and delta 0.2 (bottom), and with purities of 0.6 (left) and 0.8 (right). All simulations were run with high read depth and no differential expression. The area under the precision-recall curve for each method is shown in parentheses. In all cases and across all levels of recall, diffASE-purity (red) has higher precision than tumor sample ASE (blue). As expected, higher precision is achieved with a larger delta as well as at a higher level of purity.

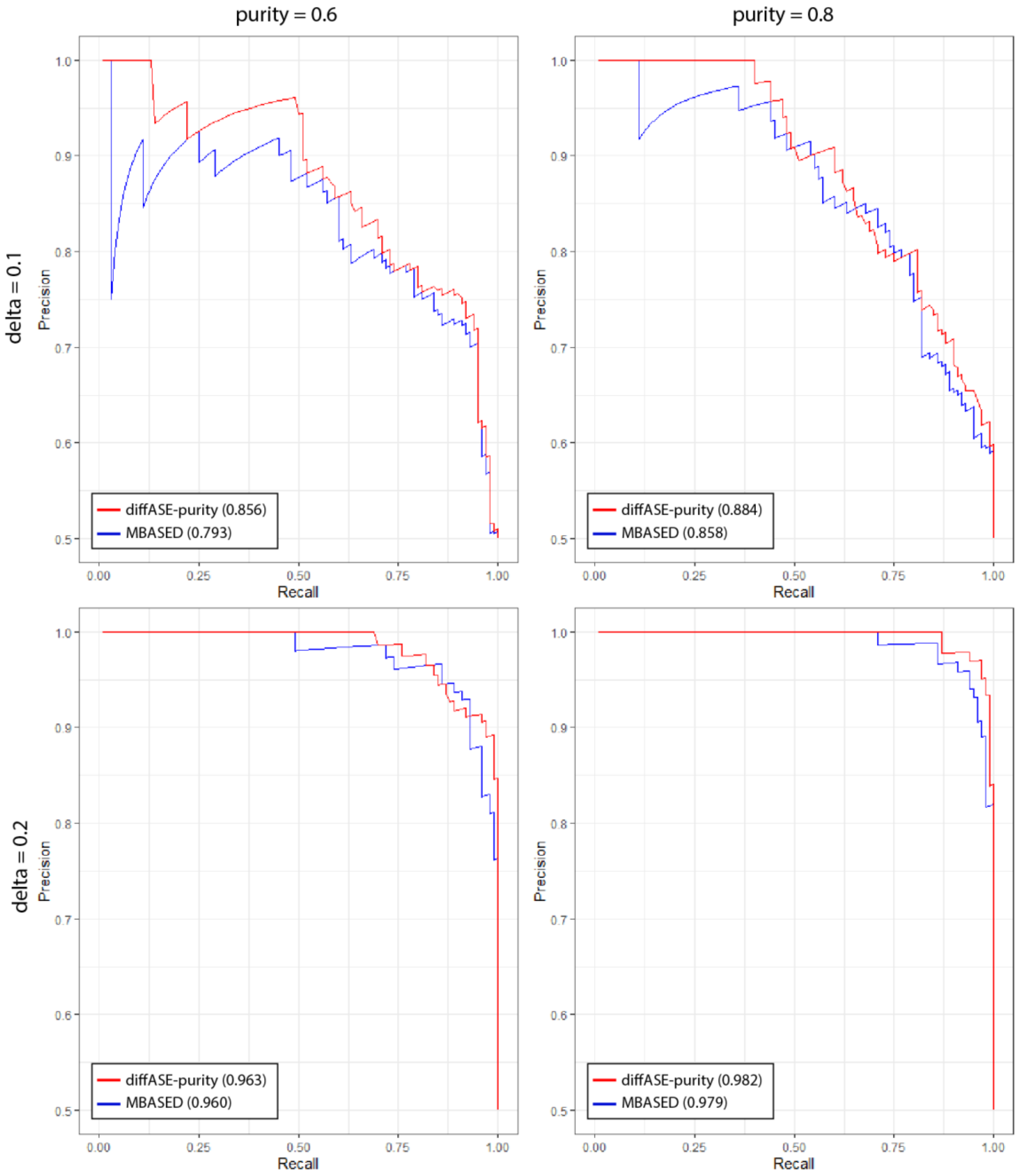


**Figure S8:** Precision-recall curves for differential ASE and two-sample MBASED. We simulated 100 negative and 100 positive sites of differential ASE as described in Section A, at delta = 0.1 (top) and delta 0.2 (bottom) with purities of 0.6 (left) and 0.8 (right). All simulations were run with high read depth and no differential expression. The area under the precision-recall curve for each method is shown in parentheses. In all cases diffASE-purity (red) has a higher area under the precision recall curve than two-sample MBASED (blue), with a consistent advantage across almost all levels of recall. We note that MBASED requires approximately 15 seconds per heterozygous site, and thus we limit comparisons to 100 sites of each type.

**Section B: CNV Effect Size**

A quantile-quantile plot (QQ-plot) of estimated differential ASE for gene-sample pairs affected by CNVs versus those that are not (**Figure S9**).

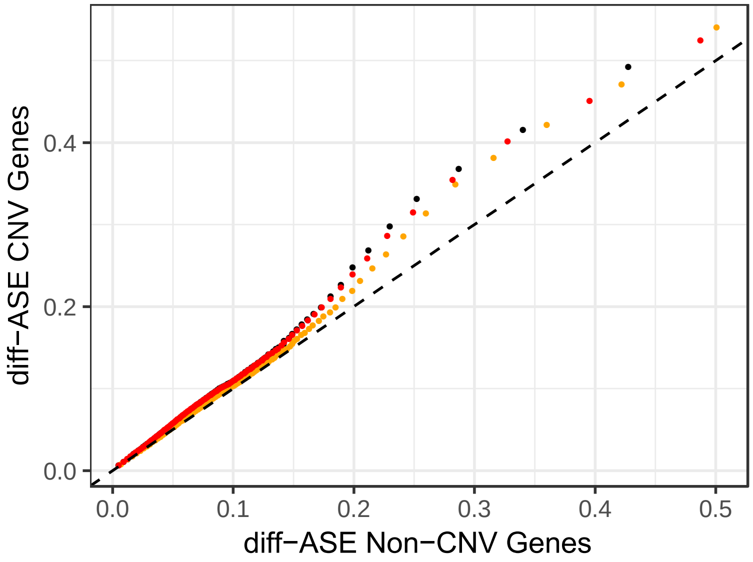


**Figure S9:** QQ-plot of differential ASE for gene-sample pairs that are affected by CNVs versus those that are not. All three models are shown with diffASE-baseline in black, diffASE-purity in orange, and diffASE-exp in red. Across all three models, gene-sample pairs with CNVs tend to have higher estimated differential ASEs.

**Section C: Genes with Recurrent Differential ASE**

**Table S1** shows genes that recurrently exhibit differential ASE. Based on the number of individuals in our study and the fraction of genes exhibiting differential ASE in each sample, we estimate that a gene must exhibit differential ASE at least six times across individuals to be statistically significant based on a Poisson binomial test and a Benjamini-Hochberg corrected FDR < 0.1.

**Table S1:** Genes with Recurrent Differential ASE

| Gene | Number of Samples Without CNV | Number of Samples With CNV |
|---|---|---|
| GSR | 20 | 3 |
| METTL7A | 18 | 0 |
| RPA1 | 17 | 2 |
| MREG | 16 | 0 |
| RABEP1 | 15 | 3 |
| NUP88 | 15 | 2 |
| MDM2 | 15 | 0 |
| CPM | 15 | 0 |
| PAICS | 14 | 0 |
| VBP1 | 13 | 0 |
| HSPA8 | 12 | 1 |
| UBQLN1 | 12 | 0 |
| MRPS16 | 12 | 0 |
| ZNF552 | 12 | 0 |
| ZDHHC20 | 12 | 2 |
| LINC00847 | 12 | 0 |
| SLC25A5 | 11 | 0 |
| PSMB2 | 11 | 0 |
| SLC25A43 | 11 | 0 |
| DNAJC9-AS1 | 11 | 0 |
| ANXA6 | 10 | 0 |
| HSPE1 | 10 | 0 |
| DYNLT3 | 10 | 0 |
| NOP14 | 10 | 0 |
| AHR | 9 | 0 |
| XIAP | 9 | 0 |
| NDUFA10 | 9 | 0 |
| CNOT1 | 9 | 3 |
| DHRS7 | 9 | 0 |
| CXCL16 | 9 | 1 |
| CHCHD1 | 9 | 0 |

| | | |
|---|---|---|
| SMIM14 | 9 | 0 |
| FGD5-AS1 | 9 | 0 |
| HSPE1-MOB4 | 9 | 0 |
| AMFR | 8 | 2 |
| C1QB | 8 | 1 |
| FUT7 | 8 | 0 |
| ABLIM1 | 8 | 0 |
| RARS | 8 | 1 |
| RPL13 | 8 | 0 |
| SHBG | 8 | 1 |
| TSPAN6 | 8 | 0 |
| UBP1 | 8 | 0 |
| ZNF75D | 8 | 0 |
| INE1 | 8 | 0 |
| ADGRG1 | 8 | 0 |
| AKR1A1 | 8 | 0 |
| ARL6IP1 | 8 | 1 |
| FBXL2 | 8 | 0 |
| HDDC2 | 8 | 0 |
| TVP23B | 8 | 1 |
| ANAPC11 | 8 | 0 |
| ARMCX3 | 8 | 0 |
| ZNF586 | 8 | 0 |
| CMC2 | 8 | 2 |
| SETD6 | 8 | 2 |
| SAT2 | 8 | 2 |
| TMEM99 | 8 | 3 |
| PGAM5 | 8 | 0 |
| NOP14-AS1 | 8 | 0 |
| LOC339260 | 8 | 3 |
| SNX19 | 8 | 2 |
| SNORD68 | 8 | 0 |
| CBR1 | 7 | 0 |
| MCM4 | 7 | 2 |
| PPIA | 7 | 0 |
| EIF2AK2 | 7 | 0 |
| RPL30 | 7 | 5 |
| RPL29 | 7 | 0 |
| RPL36AL | 7 | 0 |
| TXN | 7 | 0 |
| STK24 | 7 | 0 |

| | | |
|---|---|---|
| SLU7 | 7 | 0 |
| STRAP | 7 | 0 |
| USP22 | 7 | 2 |
| THUMPD3 | 7 | 0 |
| PDCD4 | 7 | 0 |
| MBTPS2 | 7 | 0 |
| DDX41 | 7 | 0 |
| TAF9B | 7 | 0 |
| HEATR3 | 7 | 0 |
| KIAA1143 | 7 | 0 |
| WDR48 | 7 | 0 |
| CYB5B | 7 | 3 |
| SLC38A1 | 7 | 0 |
| ASB14 | 7 | 0 |
| PLBD2 | 7 | 0 |
| CDC26 | 7 | 0 |
| C9orf142 | 7 | 0 |
| THUMPD3-AS1 | 7 | 0 |
| LIMD1-AS1 | 7 | 0 |
| MIR33B | 7 | 0 |
| UXT-AS1 | 7 | 0 |
| NDUFB2-AS1 | 7 | 0 |
| MIR1307 | 7 | 0 |
| ANXA2 | 6 | 0 |
| ARSD | 6 | 0 |
| ASS1 | 6 | 0 |
| C1QBP | 6 | 0 |
| SLC31A1 | 6 | 0 |
| SLC25A10 | 6 | 0 |
| NQO1 | 6 | 0 |
| DPAGT1 | 6 | 0 |
| GSTP1 | 6 | 0 |
| IDS | 6 | 0 |
| LAMP2 | 6 | 0 |
| SERPINA5 | 6 | 0 |
| POLR2A | 6 | 0 |
| POLR2J | 6 | 0 |
| PSMB6 | 6 | 0 |
| SAA2 | 6 | 0 |
| SLC12A2 | 6 | 0 |
| SMARCD2 | 6 | 0 |

| | | |
|---|---|---|
| UXT | 6 | 0 |
| PEX3 | 6 | 3 |
| PIR | 6 | 0 |
| SUCLG2 | 6 | 1 |
| LIMD1 | 6 | 0 |
| CWC27 | 6 | 0 |
| MAGED2 | 6 | 0 |
| GPD1L | 6 | 0 |
| SAMM50 | 6 | 0 |
| NOL11 | 6 | 0 |
| RRP7A | 6 | 0 |
| GEMIN4 | 6 | 0 |
| ARHGEF3 | 6 | 0 |
| APIP | 6 | 0 |
| TMEM14C | 6 | 0 |
| NANS | 6 | 0 |
| MPHOSPH8 | 6 | 0 |
| CWF19L1 | 6 | 0 |
| RFK | 6 | 0 |
| VPS11 | 6 | 1 |
| PERP | 6 | 3 |
| PJA1 | 6 | 0 |
| PCNX4 | 6 | 0 |
| TBC1D15 | 6 | 0 |
| APOO | 6 | 0 |
| ZNF343 | 6 | 0 |
| CERS4 | 6 | 1 |
| COASY | 6 | 0 |
| MAGT1 | 6 | 0 |
| RPAIN | 6 | 2 |
| C19orf48 | 6 | 0 |
| ZFP90 | 6 | 2 |
| TOM1L2 | 6 | 1 |
| SYCP2L | 6 | 0 |
| SUMF1 | 6 | 0 |
| RPL7L1 | 6 | 0 |
| PRSS53 | 6 | 0 |
| SNORA21 | 6 | 0 |
| LOC100133286 | 6 | 0 |
| CEBPZOS | 6 | 0 |
| SAA2-SAA4 | 6 | 0 |

| | | |
|---|---|---|
| RALY-AS1 | 6 | 0 |
| CRK | 5 | 1 |
| PSMD5 | 5 | 2 |
| PSMD12 | 5 | 2 |
| RPS6KA1 | 5 | 2 |
| DYNLT1 | 5 | 2 |
| VEZF1 | 5 | 4 |
| MPHOSPH6 | 5 | 3 |
| BPNT1 | 5 | 5 |
| PACSIN2 | 5 | 1 |
| TTLL12 | 5 | 2 |
| C22orf24 | 5 | 1 |
| RSL1D1 | 5 | 1 |
| DNAJC15 | 5 | 1 |
| AIG1 | 5 | 1 |
| IFT46 | 5 | 1 |
| ELAC2 | 5 | 2 |
| WDR82 | 5 | 1 |
| SLIRP | 5 | 1 |
| HSDL1 | 5 | 3 |
| BTBD6 | 5 | 1 |
| LRRC75A-AS1 | 5 | 2 |
| ESR1 | 5 | 2 |
| ACAT1 | 4 | 2 |
| ASAH1 | 4 | 6 |
| FUCA2 | 4 | 4 |
| STX8 | 4 | 2 |
| SNW1 | 4 | 2 |
| MRTO4 | 4 | 2 |
| RCBTB1 | 4 | 3 |
| LDOC1L | 4 | 2 |
| LRRC75A | 4 | 2 |
| PARP4 | 3 | 3 |
| EPHX2 | 3 | 3 |
| NENF | 3 | 5 |
| UTP18 | 3 | 3 |
| PHF11 | 3 | 4 |
| CCDC25 | 3 | 4 |
| S100A14 | 3 | 3 |
| RNF26 | 3 | 3 |
| TATDN1 | 2 | 4 |

| | | |
|---|---|---|
| PRUNE | 2 | 6 |
| PPP2R5A | 1 | 5 |
| FDFT1 | 0 | 7 |
| NSL1 | 0 | 6 |
| KMO | 0 | 6 |
| HSBP1 | 0 | 6 |
| COPA | 0 | 6 |
| NAT1 | 0 | 6 |

**Section D: Gene Set Enrichment Analysis**

Full lists of Reactome pathways (**Table S2**) and disease ontologies (**Table S3**) found significant by Gene Set Enrichment Analysis on the full set of genes ordered by recurrence (as given in Table S1), and sorted by Normalized Enrichment Score (NES) (Yu & He, 2016) .

**Table S2:** Reactome pathways

| ID | Description | Set Size | NES | *p*-value |
|---|---|---|---|---|
| 69541 | Stabilization of p53 | 30 | 1.79 | 1.01E-03 |
| 5578749 | Transcriptional regulation by small RNAs | 26 | 1.77 | 5.10E-03 |
| 72172 | mRNA Splicing | 50 | 1.76 | 9.99E-04 |
| 72163 | mRNA Splicing - Major Pathway | 50 | 1.76 | 9.99E-04 |
| 1799339 | SRP-dependent cotranslational protein targeting to membrane | 68 | 1.76 | 2.00E-03 |
| 69620 | Cell Cycle Checkpoints | 61 | 1.74 | 9.99E-04 |
| 69615 | G1/S DNA Damage Checkpoints | 31 | 1.73 | 1.01E-03 |
| 69563 | p53-Dependent G1 DNA Damage Response | 31 | 1.73 | 1.01E-03 |
| 69580 | p53-Dependent G1/S DNA damage checkpoint | 31 | 1.73 | 1.01E-03 |
| 72203 | Processing of Capped Intron-Containing Pre-mRNA | 53 | 1.73 | 9.99E-04 |
| 167161 | HIV Transcription Initiation | 16 | 1.72 | 3.19E-03 |
| 167162 | RNA Polymerase II HIV Promoter Escape | 16 | 1.72 | 3.19E-03 |
| 73776 | RNA Polymerase II Promoter Escape | 16 | 1.72 | 3.19E-03 |
| 75953 | RNA Polymerase II Transcription Initiation | 16 | 1.72 | 3.19E-03 |
| 76042 | RNA Polymerase II Transcription Initiation And Promoter Clearance | 16 | 1.72 | 3.19E-03 |
| 73779 | RNA Polymerase II Transcription Pre-Initiation And Promoter Opening | 16 | 1.72 | 3.19E-03 |
| 73857 | RNA Polymerase II Transcription | 43 | 1.71 | 1.00E-03 |
| 179419 | APC:Cdc20 mediated degradation of cell cycle proteins prior to satisfation of the cell cycle checkpoint | 39 | 1.71 | 1.01E-03 |
| 174184 | Cdc20:Phospho-APC/C mediated degradation of Cyclin A | 39 | 1.71 | 1.01E-03 |
| 69002 | DNA Replication Pre-Initiation | 40 | 1.71 | 2.01E-03 |
| 68874 | M/G1 Transition | 40 | 1.71 | 2.01E-03 |
| 450408 | AUF1 (hnRNP D0) destabilizes mRNA | 32 | 1.71 | 3.02E-03 |
| 168254 | Influenza Infection | 75 | 1.70 | 9.99E-04 |
| 499943 | Synthesis and interconversion of nucleotide di- and triphosphates | 13 | 1.70 | 7.45E-03 |
| 168273 | Influenza Viral RNA Transcription and Replication | 71 | 1.69 | 2.00E-03 |
| 73885 | Nucleotide Excision Repair | 22 | 1.69 | 4.14E-03 |
| 73937 | Transcription-coupled NER (TC-NER) | 22 | 1.69 | 4.14E-03 |
| 3299685 | Detoxification of Reactive Oxygen Species | 17 | 1.69 | 6.34E-03 |
| 176814 | Activation of APC/C and APC/C:Cdc20 mediated degradation of mitotic proteins | 41 | 1.68 | 1.01E-03 |
| 176409 | APC/C:Cdc20 mediated degradation of mitotic proteins | 41 | 1.68 | 1.01E-03 |
| 72165 | mRNA Splicing - Minor Pathway | 21 | 1.68 | 8.26E-03 |
| 350562 | Regulation of ornithine decarboxylase (ODC) | 29 | 1.67 | 7.11E-03 |

| | | | | |
|---|---|---|---|---|
| 68962 | Activation of the pre-replicative complex | 11 | 1.67 | 6.70E-03 |
| 211000 | Regulatory RNA pathways | 31 | 1.67 | 9.13E-03 |
| 174084 | Autodegradation of Cdh1 by Cdh1:APC/C | 38 | 1.67 | 4.02E-03 |
| 72766 | Translation | 82 | 1.67 | 9.99E-04 |
| 174154 | APC/C:Cdc20 mediated degradation of Securin | 38 | 1.66 | 4.03E-03 |
| 72764 | Eukaryotic Translation Termination | 55 | 1.66 | 2.00E-03 |
| 192823 | Viral mRNA Translation | 55 | 1.66 | 2.00E-03 |
| 168255 | Influenza Life Cycle | 73 | 1.66 | 2.00E-03 |
| 156842 | Eukaryotic Translation Elongation | 57 | 1.65 | 3.00E-03 |
| 156902 | Peptide chain elongation | 57 | 1.65 | 3.00E-03 |
| 4641257 | degradation of AXIN | 31 | 1.64 | 6.06E-03 |
| 179409 | APC-Cdc20 mediated degradation of Nek2A | 12 | 1.64 | 4.37E-03 |
| 71291 | Metabolism of amino acids and derivatives | 85 | 1.64 | 9.99E-04 |
| 674695 | RNA Polymerase II Pre-transcription Events | 25 | 1.63 | 8.16E-03 |
| 975956 | Nonsense Mediated Decay (NMD) independent of the Exon Junction Complex (EJC) | 57 | 1.63 | 3.00E-03 |
| 1428517 | The citric acid (TCA) cycle and respiratory electron transport | 68 | 1.63 | 3.00E-03 |
| 174178 | APC/C:Cdh1 mediated degradation of Cdc20 and other APC/C:Cdh1 targeted proteins in late mitosis/early G1 | 40 | 1.62 | 5.02E-03 |
| 176408 | Regulation of APC/C activators between G1/S and early anaphase | 45 | 1.62 | 4.01E-03 |
| 174143 | APC/C-mediated degradation of cell cycle proteins | 47 | 1.62 | 4.01E-03 |
| 453276 | Regulation of mitotic cell cycle | 47 | 1.62 | 4.01E-03 |
| 187577 | SCF(Skp2)-mediated degradation of p27/p21 | 32 | 1.61 | 9.05E-03 |
| 162599 | Late Phase of HIV Life Cycle | 62 | 1.61 | 9.99E-04 |
| 72689 | Formation of a pool of free 40S subunits | 60 | 1.61 | 3.00E-03 |
| 349425 | Autodegradation of the E3 ubiquitin ligase COP1 | 29 | 1.61 | 1.01E-02 |
| 69610 | p53-Independent DNA Damage Response | 29 | 1.61 | 1.01E-02 |
| 69613 | p53-Independent G1/S DNA damage checkpoint | 29 | 1.61 | 1.01E-02 |
| 69601 | Ubiquitin Mediated Degradation of Phosphorylated Cdc25A | 29 | 1.61 | 1.01E-02 |
| 75815 | Ubiquitin-dependent degradation of Cyclin D | 29 | 1.61 | 1.01E-02 |
| 69229 | Ubiquitin-dependent degradation of Cyclin D1 | 29 | 1.61 | 1.01E-02 |
| 68867 | Assembly of the pre-replicative complex | 35 | 1.60 | 5.03E-03 |
| 2029480 | Fcgamma receptor (FCGR) dependent phagocytosis | 30 | 1.60 | 9.10E-03 |
| 611105 | Respiratory electron transport | 41 | 1.60 | 2.00E-03 |
| 5610783 | Degradation of GLI2 by the proteasome | 34 | 1.60 | 6.01E-03 |
| 5663205 | Infectious disease | 185 | 1.60 | 9.99E-04 |
| 195253 | Degradation of beta-catenin by the destruction complex | 38 | 1.59 | 5.03E-03 |
| 5607761 | Dectin-1 mediated noncanonical NF-kB signaling | 35 | 1.59 | 6.03E-03 |
| 69017 | CDK-mediated phosphorylation and removal of Cdc6 | 30 | 1.59 | 1.01E-02 |
| 157279 | 3' -UTR-mediated translational regulation | 64 | 1.58 | 3.00E-03 |
| 156827 | L13a-mediated translational silencing of Ceruloplasmin expression | 64 | 1.58 | 3.00E-03 |
| 1169091 | Activation of NF-kappaB in B cells | 38 | 1.58 | 5.02E-03 |

| | | | | |
|---|---|---|---|---|
| 163200 | Respiratory electron transport, ATP synthesis by chemiosmotic coupling, and heat production by uncoupling proteins. | 45 | 1.58 | 3.01E-03 |
| 5610785 | GLI3 is processed to GLI3R by the proteasome | 35 | 1.57 | 6.01E-03 |
| 72706 | GTP hydrolysis and joining of the 60S ribosomal subunit | 64 | 1.57 | 4.00E-03 |
| 109581 | Apoptosis | 70 | 1.57 | 4.00E-03 |
| 5357801 | Programmed Cell Death | 70 | 1.57 | 4.00E-03 |
| 69206 | G1/S Transition | 50 | 1.56 | 5.01E-03 |
| 162587 | HIV Life Cycle | 69 | 1.56 | 2.00E-03 |
| 15869 | Metabolism of nucleotides | 37 | 1.56 | 5.01E-03 |
| 5607764 | CLEC7A (Dectin-1) signaling | 51 | 1.56 | 4.00E-03 |
| 72737 | Cap-dependent Translation Initiation | 67 | 1.56 | 4.00E-03 |
| 72613 | Eukaryotic Translation Initiation | 67 | 1.56 | 4.00E-03 |
| 69306 | DNA Replication | 48 | 1.56 | 5.01E-03 |
| 69239 | Synthesis of DNA | 48 | 1.56 | 5.01E-03 |
| 5621481 | C-type lectin receptors (CLRs) | 60 | 1.55 | 5.00E-03 |
| 1168372 | Downstream signaling events of B Cell Receptor (BCR) | 68 | 1.54 | 3.00E-03 |
| 450531 | Regulation of mRNA stability by proteins that bind AU-rich elements | 46 | 1.54 | 6.00E-03 |
| 453279 | Mitotic G1-G1/S phases | 54 | 1.54 | 7.00E-03 |
| 68949 | Orc1 removal from chromatin | 37 | 1.53 | 7.04E-03 |
| 69304 | Regulation of DNA replication | 37 | 1.53 | 7.04E-03 |
| 69300 | Removal of licensing factors from origins | 37 | 1.53 | 7.04E-03 |
| 69052 | Switching of origins to a post-replicative state | 37 | 1.53 | 7.04E-03 |
| 3858494 | beta-catenin independent WNT signaling | 49 | 1.52 | 8.00E-03 |
| 162906 | HIV Infection | 112 | 1.51 | 9.99E-04 |
| 69242 | S Phase | 55 | 1.50 | 8.00E-03 |
| 2262752 | Cellular responses to stress | 112 | 1.49 | 2.00E-03 |
| 983705 | Signaling by the B Cell Receptor (BCR) | 77 | 1.48 | 5.00E-03 |
| 162909 | Host Interactions of HIV factors | 71 | 1.48 | 5.00E-03 |
| 1643685 | Disease | 282 | 1.48 | 9.99E-04 |
| 187037 | NGF signalling via TRKA from the plasma membrane | 64 | 1.47 | 8.99E-03 |
| 392499 | Metabolism of proteins | 296 | 1.47 | 9.99E-04 |
| 168249 | Innate Immune System | 188 | 1.46 | 9.99E-04 |
| 74160 | Gene Expression | 377 | 1.39 | 9.99E-04 |
| 597592 | Post-translational protein modification | 134 | 1.38 | 9.99E-04 |
| 168256 | Immune System | 307 | 1.36 | 9.99E-04 |
| 1280218 | Adaptive Immune System | 140 | 1.34 | 6.99E-03 |

**Table S3:** Disease Ontologies

| ID | Description | Set Size | NES | *p*-value |
|---|---|---|---|---|
| DOID:8567 | Hodgkin's lymphoma | 17 | 1.86 | 1.04E-03 |
| DOID:3748 | esophagus squamous cell carcinoma | 18 | 1.81 | 3.17E-03 |
| DOID:9261 | nasopharynx carcinoma | 13 | 1.77 | 3.29E-03 |
| DOID:0060119 | pharynx cancer | 14 | 1.73 | 5.41E-03 |
| DOID:0060058 | Lymphoma | 29 | 1.73 | 1.01E-03 |
| DOID:10155 | intestinal cancer | 117 | 1.69 | 9.99E-04 |
| DOID:219 | colon cancer | 111 | 1.68 | 9.99E-04 |
| DOID:9256 | colorectal cancer | 111 | 1.68 | 9.99E-04 |
| DOID:5672 | large intestine cancer | 112 | 1.67 | 9.99E-04 |
| DOID:4079 | heart valve disease | 13 | 1.67 | 1.08E-02 |
| DOID:12603 | acute leukemia | 30 | 1.65 | 4.04E-03 |
| DOID:4989 | Pancreatitis | 29 | 1.62 | 6.08E-03 |
| DOID:26 | pancreas disease | 40 | 1.60 | 6.00E-03 |
| DOID:1520 | colon carcinoma | 64 | 1.60 | 2.00E-03 |
| DOID:10283 | prostate cancer | 160 | 1.58 | 9.99E-04 |
| DOID:6000 | congestive heart failure | 58 | 1.58 | 3.00E-03 |
| DOID:3856 | male reproductive organ cancer | 161 | 1.58 | 9.99E-04 |
| DOID:1040 | chronic lymphocytic leukemia | 58 | 1.55 | 4.00E-03 |
| DOID:193 | reproductive organ cancer | 264 | 1.55 | 9.99E-04 |
| DOID:1749 | squamous cell carcinoma | 252 | 1.54 | 9.99E-04 |
| DOID:0060056 | hypersensitivity reaction disease | 191 | 1.54 | 9.99E-04 |
| DOID:1037 | lymphoblastic leukemia | 127 | 1.53 | 9.99E-04 |
| DOID:326 | Ischemia | 64 | 1.53 | 5.00E-03 |
| DOID:229 | female reproductive system disease | 46 | 1.51 | 8.01E-03 |
| DOID:417 | hypersensitivity reaction type II disease | 163 | 1.51 | 9.99E-04 |
| DOID:936 | brain disease | 101 | 1.51 | 9.99E-04 |
| DOID:3119 | gastrointestinal system cancer | 438 | 1.50 | 9.99E-04 |
| DOID:1883 | hepatitis C | 71 | 1.49 | 4.00E-03 |
| DOID:0060085 | organ system benign neoplasm | 90 | 1.48 | 4.00E-03 |
| DOID:305 | Carcinoma | 292 | 1.48 | 9.99E-04 |
| DOID:1324 | lung cancer | 202 | 1.48 | 9.99E-04 |
| DOID:28 | endocrine system disease | 95 | 1.48 | 3.00E-03 |
| DOID:15 | reproductive system disease | 87 | 1.47 | 5.99E-03 |
| DOID:0050615 | respiratory system cancer | 206 | 1.47 | 9.99E-04 |
| DOID:0060072 | benign neoplasm | 176 | 1.47 | 9.99E-04 |
| DOID:3963 | thyroid carcinoma | 74 | 1.46 | 6.99E-03 |
| DOID:0060083 | immune system cancer | 360 | 1.46 | 9.99E-04 |
| DOID:4074 | pancreas adenocarcinoma | 61 | 1.45 | 1.10E-02 |
| DOID:2531 | hematologic cancer | 359 | 1.45 | 9.99E-04 |
| DOID:10534 | stomach cancer | 89 | 1.45 | 6.99E-03 |

| | | | | |
|---|---|---|---|---|
| DOID:170 | endocrine gland cancer | 169 | 1.44 | 2.00E-03 |
| DOID:17 | musculoskeletal system disease | 359 | 1.44 | 9.99E-04 |
| DOID:4766 | Embryoma | 119 | 1.44 | 3.00E-03 |
| DOID:2914 | immune system disease | 307 | 1.44 | 9.99E-04 |
| DOID:65 | connective tissue disease | 277 | 1.44 | 9.99E-04 |
| DOID:1781 | thyroid cancer | 76 | 1.43 | 8.99E-03 |
| DOID:423 | Myopathy | 132 | 1.43 | 2.00E-03 |
| DOID:66 | muscle tissue disease | 132 | 1.43 | 2.00E-03 |
| DOID:1240 | Leukemia | 321 | 1.43 | 9.99E-04 |
| DOID:0060084 | cell type benign neoplasm | 141 | 1.43 | 4.00E-03 |
| DOID:850 | lung disease | 124 | 1.42 | 5.00E-03 |
| DOID:3996 | urinary system cancer | 180 | 1.42 | 2.00E-03 |
| DOID:3905 | lung carcinoma | 182 | 1.42 | 9.99E-04 |
| DOID:688 | embryonal cancer | 123 | 1.42 | 3.00E-03 |
| DOID:3908 | non-small cell lung carcinoma | 157 | 1.42 | 9.99E-04 |
| DOID:0080000 | muscular disease | 134 | 1.42 | 9.99E-04 |
| DOID:2621 | autonomic nervous system neoplasm | 120 | 1.41 | 5.00E-03 |
| DOID:769 | Neuroblastoma | 120 | 1.41 | 5.00E-03 |
| DOID:374 | nutrition disease | 85 | 1.41 | 7.99E-03 |
| DOID:120 | female reproductive organ cancer | 148 | 1.41 | 3.00E-03 |
| DOID:4451 | renal carcinoma | 140 | 1.41 | 4.00E-03 |
| DOID:3118 | hepatobiliary disease | 191 | 1.41 | 2.00E-03 |
| DOID:657 | Adenoma | 99 | 1.40 | 7.99E-03 |
| DOID:263 | kidney cancer | 158 | 1.40 | 4.00E-03 |
| DOID:409 | liver disease | 181 | 1.40 | 2.00E-03 |
| DOID:1909 | Melanoma | 233 | 1.40 | 9.99E-04 |
| DOID:2237 | Hepatitis | 126 | 1.40 | 6.99E-03 |
| DOID:0050161 | lower respiratory tract disease | 128 | 1.40 | 6.99E-03 |
| DOID:331 | central nervous system disease | 349 | 1.40 | 9.99E-04 |
| DOID:934 | viral infectious disease | 230 | 1.40 | 9.99E-04 |
| DOID:201 | connective tissue cancer | 129 | 1.40 | 5.00E-03 |
| DOID:3571 | liver cancer | 311 | 1.40 | 9.99E-04 |
| DOID:684 | hepatocellular carcinoma | 311 | 1.40 | 9.99E-04 |
| DOID:686 | liver carcinoma | 311 | 1.40 | 9.99E-04 |
| DOID:0060100 | musculoskeletal system cancer | 150 | 1.39 | 2.00E-03 |
| DOID:0080001 | bone disease | 232 | 1.39 | 9.99E-04 |
| DOID:0050117 | disease by infectious agent | 278 | 1.38 | 9.99E-04 |
| DOID:1579 | respiratory system disease | 134 | 1.38 | 8.99E-03 |
| DOID:2994 | germ cell cancer | 134 | 1.38 | 5.00E-03 |
| DOID:10652 | Alzheimer's disease | 136 | 1.38 | 5.00E-03 |
| DOID:680 | Tauopathy | 136 | 1.38 | 5.00E-03 |
| DOID:1192 | peripheral nervous system neoplasm | 128 | 1.38 | 8.99E-03 |
| DOID:3093 | nervous system cancer | 191 | 1.37 | 9.99E-04 |

| | | | | |
|---|---|---|---|---|
| DOID:1793 | pancreatic cancer | 110 | 1.37 | 8.99E-03 |
| DOID:4450 | renal cell carcinoma | 128 | 1.36 | 5.99E-03 |
| DOID:3342 | bone inflammation disease | 193 | 1.36 | 9.99E-04 |
| DOID:114 | heart disease | 145 | 1.36 | 9.99E-03 |
| DOID:863 | nervous system disease | 470 | 1.34 | 9.99E-04 |
| DOID:1612 | breast cancer | 191 | 1.34 | 6.99E-03 |
| DOID:5093 | thoracic cancer | 191 | 1.34 | 6.99E-03 |
| DOID:1287 | cardiovascular system disease | 359 | 1.33 | 2.00E-03 |
| DOID:77 | gastrointestinal system disease | 255 | 1.33 | 4.00E-03 |
| DOID:848 | Arthritis | 183 | 1.33 | 3.00E-03 |
| DOID:178 | vascular disease | 293 | 1.32 | 2.00E-03 |
| DOID:150 | disease of mental health | 331 | 1.32 | 3.00E-03 |
| DOID:1289 | neurodegenerative disease | 262 | 1.31 | 5.99E-03 |
| DOID:3459 | breast carcinoma | 145 | 1.30 | 1.10E-02 |
| DOID:1561 | cognitive disorder | 250 | 1.30 | 5.99E-03 |
| DOID:0050828 | artery disease | 244 | 1.29 | 1.10E-02 |
| DOID:225 | Syndrome | 471 | 1.28 | 2.00E-03 |

**Section E: Enrichment for Mutation Types Using Tumor Sample ASE**

Among gene-sample pairs with significant ASE in the tumor sample, we calculate the enrichment for different mutation types. Across different mutation types, enrichments are modest, even for genes with nonsense mutations. When comparing enrichments obtained in gene-sample pairs identified as significant via tumor sample ASE or differential ASE, enrichments for coding mutations are similar. However, for noncoding mutations, only 3'UTR mutations had a very small but significant enrichment with tumor sample ASE (**Table S4**, significant categories are marked with an asterisk at hypergeometric *p*-value <05). This indicates that, unlike differential ASE, tumor sample ASE is not able to detect the effects of *cis* noncoding mutations.

**Table S4:** Enrichment for Mutation Types Using Tumor Sample ASE

| Mutation Type | Enrichment | Number of Significant Mutations |
|---|---|---|
| Nonsense | 1.09* | 523 |
| Nonsense <50bp from 3'UTR | 1.27 | 17 |
| Missense | 1.08* | 5,588 |
| Silent | 1.10* | 1,101 |
| Promoter | 0.95 | 229 |
| Annotated Regulatory Region in Promoter | 0.95 | 92 |
| 5'UTR | 1.05 | 160 |
| 3'UTR | 1.14* | 736 |
| 3'Flank | 1.04 | 234 |
| Intron | 1.00 | 7,505 |
| Unmutated | 0.997 | - |

**Section F: Mutation Counts**

**Table S5** is a summary of the total numbers of each mutation type we observed in TCGA exome (WES) data across the 91 individuals we studied and the 46 remaining samples after removing those whose correlations with immune gene profiles are significant according to Cibersort (Newman et al., 2015; TCGA Research Network).

One of the most powerful features of differential ASE is that it serves as a filter for which *cis* noncoding mutations have the potential to cause dysregulation. **Table S6** shows the total numbers of each noncoding mutation type in locations that are evolutionarily conserved (Pollard, Hubisz, Rosenbloom, & Siepel, 2010), then of those, how many are *cis* to genes with significant differential ASE (at FDR < 0.1), and then finally, of those, which are in regulatory regions as annotated by ORegAnno (Lesurf et al., 2016) .

**Table S5:** Mutation Counts by Type in TCGA

| Mutation Type | Mutation Count | Cibersort Filtered Count |
|---|---|---|
| Nonsense | 4,320 | 2,261 |
| Missense | 71,231 | 37,555 |
| Silent | 28,197 | 14,672 |
| Promoter | 12,026 | 5,042 |
| 5'UTR | 3,704 | 1,882 |
| 3'UTR | 12,778 | 6,732 |
| 3'Flank | 17,337 | 7,517 |
| Intron | 257,570 | 117,903 |

**Table S6:** Mutation Counts Filtered By Differential ASE

| Mutation Type | Conserved Mutation Count | Significant Differential ASE (FDR < 0.1) | In Annotated Regulatory Region |
|---|---|---|---|
| Promoter | 583 | 19 | 9 |
| 5'UTR | 389 | 20 | 15 |
| 3'UTR | 1,678 | 97 | 15 |
| 3'Flank | 684 | 23 | 3 |
| Intron | 20,066 | 716 | 105 |

**Section G: Full List of Potential Functional Noncoding Mutations**
This section contains 5 tables by type of mutations *cis* to genes exhibiting significant differential ASE (FDR < 0.1) that occur within annotated regulatory regions. **Table S7** shows promoter mutations, **Table S8** shows 5'UTR mutations, **Table S9** shows 3'UTR mutations, **Table S10** shows 3'Flank mutations, and **Table S11** shows Intron mutations. In cases where two different mutations were observed in the same individual *cis* to a single gene exhibiting differential ASE, those locations are listed without renaming the gene, as we cannot distinguish between the contributions of the two mutations.

**Table S7:** Promoter Mutations

| Gene | Chromosome | Location | Reference Allele | Tumor Allele |
|---|---|---|---|---|
| ATP6V0E2 | chr7 | 149873827 | G | T |
| GNB1 | chr1 | 1892263 | A | C |
| | | 1892276 | A | C |
| SRPK2 | chr7 | 105389327 | G | A |
| S100A11 | chr1 | 152041960 | A | T |
| PNKD | chr2 | 218270466 | C | A |
| PPP6R3 | chr11 | 68458148 | A | G |
| PEF1 | chr1 | 31645237 | T | G |
| CUEDC2 | chr10 | 102436521 | G | C |

**Table S8:** 5'UTR Mutations

| Gene | Chromosome | Location | Reference Allele | Tumor Allele |
|---|---|---|---|---|
| PGRMC1 | chrX | 119236356 | C | A |
| UBE2J1 | chr6 | 89352574 | G | T |
| RPA2 | chr1 | 27914489 | G | T |
| ZMAT2 | chr5 | 140700411 | A | C |
| RHOB | chr2 | 20447402 | C | T |
| C14orf166 | chr14 | 51989638 | C | T |
| GADD45B | chr19 | 2476343 | G | T |
| ZFP36L1 | chr14 | 68793124 | G | A |
| AREL1 | chr14 | 74713092 | A | C |
| IL1RN | chr2 | 113127611 | G | T |
| ST7 | chr7 | 116953433 | G | T |
| HPS4 | chr22 | 26481782 | A | G |
| WDR48 | chr3 | 39052005 | G | T |
| TAF9B | chrX | 78139616 | T | C |
| COASY | chr17 | 42562291 | T | A |
| ZNF552 | chr19 | 57814766 | G | A |
| CDKN1B | chr12 | 12717761 | T | C |

**Table S9:** 3’UTR Mutations

| Gene | Chromosome | Location | Reference Allele | Tumor Allele |
|---|---|---|---|---|
| PAXX | chr9 | 136993881 | A | C |
| PAXX | chr9 | 136993900 | A | C |
| ANXA6 | chr5 | 151101356 | C | A |
| | | 151101384 | C | A |
| ZDHHC11 | chr5 | 796106 | G | A |
| HYOU1 | chr11 | 119044760 | G | T |
| INPPL1 | chr11 | 72238722 | G | A |
| SRP54 | chr14 | 35029533 | G | T |
| CLINT1 | chr5 | 157787068 | C | A |
| DGCR2 | chr22 | 19037961 | C | A |
| TBL1X | chrX | 9716285 | A | C |
| VSIG4 | chrX | 66022148 | G | T |
| MYL6 | chr12 | 56160681 | T | C |

**Table S10:** 3’Flank Mutations

| Gene | Chromosome | Location | Reference Allele | Tumor Allele |
|---|---|---|---|---|
| SP1 | chr12 | 53413154 | C | A |
| GBP2 | chr1 | 89104347 | C | T |
| STRN3 | chr14 | 30894862 | G | T |

**Table S11:** Intron Mutations:

| Gene | Chromosome | Location | Reference Allele | Tumor Allele |
|---|---|---|---|---|
| GINS2 | chr16 | 85678680 | G | T |
| IPO7 | chr11 | 9429477 | C | A |
| AKT1 | chr14 | 104780333 | G | T |
| USP24 | chr1 | 55172364 | A | T |
| TRUB2 | chr9 | 128322278 | G | T |
| CUTC | chr10 | 99754957 | A | T |
| SLC12A2 | chr5 | 128149887 | A | G |
| TGIF1 | chr18 | 3456220 | G | T |
| GLYR1 | chr16 | 4832929 | A | G |
| RUNX1 | chr21 | 34834648 | G | T |
| SLFN5 | chr17 | 35256951 | C | A |
| GALNS | chr16 | 88824198 | G | A |
| | | 88839558 | G | A |
| ARFGAP3 | chr22 | 42801028 | A | G |
| HDAC7 | chr12 | 47802360 | T | C |
| TRIB3 | chr20 | 363435 | T | C |
| TMTC4 | chr13 | 100673796 | C | T |
| MED7 | chr5 | 157148842 | A | C |

| | | | | |
|---|---|---|---|---|
| SLK | chr10 | 103967936 | A | C |
| SLC24A3 | chr20 | 19621398 | G | A |
| PIP5K1C | chr19 | 3643564 | C | A |
| NFKBIE | chr6 | 44264337 | A | G |
| NARFL | chr16 | 739902 | G | T |
| MCM3 | chr6 | 52272479 | T | C |
| EXOSC10 | chr1 | 11069505 | C | T |
| MAN1B1 | chr9 | 137089566 | G | T |
| GBP5 | chr1 | 89262196 | A | G |
| OCIAD1 | chr4 | 48831502 | A | G |
| PRKG1 | chr10 | 51074535 | G | A |
| PPP1R1B | chr17 | 39629910 | G | T |
| MMACHC | chr1 | 45508662 | C | T |
| ANK3 | chr10 | 60042941 | G | T |
| DNAJA4 | chr15 | 78266515 | C | T |
| ARHGAP44 | chr17 | 12949591 | T | G |
| CAPN1 | chr11 | 65187163 | G | T |
| IRF2 | chr4 | 184473956 | G | T |
| PPIH | chr1 | 42658765 | T | G |
| GPD1L | chr3 | 32140632 | T | C |
| NDUFB1 | chr14 | 92116489 | A | G |
| SIPA1L3 | chr19 | 38003864 | G | A |
| HYOU1 | chr11 | 119052582 | T | G |
| | | 119052618 | A | C |
| FCHO2 | chr5 | 72989275 | G | T |
| SLC25A43 | chrX | 119406820 | C | A |
| C2orf69 | chr2 | 199911830 | C | T |
| ERBB3 | chr12 | 56087661 | C | A |
| BRMS1 | chr11 | 66338226 | A | C |
| NOC2L | chr1 | 959175 | C | A |
| MRPL13 | chr8 | 120445037 | A | C |
| OLA1 | chr2 | 174172369 | C | T |
| HMBS | chr11 | 119089614 | T | G |
| | | 119089618 | T | G |
| HMGA1 | chr6 | 34243314 | C | G |
| INSR | chr19 | 7153383 | C | T |
| ITGB4 | chr17 | 75737295 | T | G |
| NDFIP2 | chr13 | 79533279 | T | A |
| HAUS6 | chr9 | 19078126 | A | G |
| | | 19093019 | C | G |
| EXD2 | chr14 | 69237494 | A | C |
| EDEM2 | chr20 | 35145081 | C | G |
| ERBB2IP | chr5 | 65992603 | A | G |
| NCLN | chr19 | 3192699 | G | T |

| | | | | |
|---|---|---|---|---|
| PTPRG | chr3 | 62201592 | T | G |
| C8orf33 | chr8 | 145053035 | A | C |
| | | 145052992 | G | T |
| SMARCD2 | chr17 | 63834585 | A | G |
| BNIP2 | chr15 | 59680585 | G | C |
| ST14 | chr11 | 130188487 | T | G |
| UBE2D3 | chr4 | 102801622 | T | A |
| PQLC1 | chr18 | 79943223 | A | C |
| CPNE3 | chr8 | 86558210 | A | G |
| OCIAD1 | chr4 | 48831502 | A | G |
| TSSC1 | chr2 | 3294313 | C | G |
| ANK3 | chr10 | 60187148 | A | T |
| HNRNPH3 | chr10 | 68338066 | T | G |
| AMBRA1 | chr11 | 46508518 | A | G |
| EIF2AK2 | chr2 | 37156023 | G | A |
| WNK1 | chr12 | 886370 | A | T |
| SRD5A3-AS1 | chr4 | 55381628 | C | A |
| STX8 | chr17 | 9256768 | G | A |
| PSD3 | chr8 | 18980577 | A | C |
| SH3GL1 | chr19 | 4367071 | T | C |
| PHPT1 | chr9 | 136850208 | C | T |
| TLDC1 | chr16 | 84514394 | T | C |
| | | 84514424 | C | G |
| NANS | chr9 | 98072097 | C | G |
| RPS2 | chr16 | 1963402 | G | A |
| CD63 | chr12 | 55727138 | G | T |
| HBS1L | chr6 | 135004579 | A | G |
| ITPR3 | chr6 | 33658548 | T | G |
| MED31 | chr17 | 6648300 | T | C |
| RBM18 | chr9 | 122264385 | G | A |
| UBQLN1 | chr9 | 83666798 | A | T |
| UBR3 | chr2 | 169932798 | A | C |
| STK39 | chr2 | 167955846 | C | T |
| SLC26A5 | chr7 | 103392552 | G | A |
| AP3S2 | chr15 | 89888459 | T | G |
| ZBTB49 | chr4 | 4313226 | A | C |
| PYGL | chr14 | 50920658 | A | C |
| PLEKHA8 | chr7 | 30093752 | A | G |
| DAPK1 | chr9 | 87684236 | C | T |
| ATP1A1 | chr1 | 116374149 | A | C |
| FXR2 | chr17 | 7601287 | G | T |
| EXOC4 | chr7 | 134054948 | A | T |
| ERN1 | chr17 | 64074967 | A | G |

| | | | | |
|---|---|---|---|---|
| FAM178A | chr10 | 100913306 | G | T |
| ANKRD30A | chr10 | 37197348 | A | T |

**Section H: Potential Functional Promoter Mutations from Whole-Genome Sequencing**

Four of the samples considered additionally had whole-genome sequencing (WGS) data. We looked for mutations within 2000bp of transcription start sites of genes with significant differential ASE (FDR < 0.01) that occur within annotated regulatory regions and found 5 additional potential functional mutations listed in **Table S12**.

**Table S12:** Additional Mutations in Promoters in WGS

| Gene | Chromosome | Location | Reference Allele | Tumor Allele |
|---|---|---|---|---|
| KIF1B | chr1 | 10230988 | T | G |
| TNFAIP8 | chr5 | 119267282 | A | C |
| PNN | chr14 | 39174277 | A | G |
| FCF1 | chr14 | 74713160 | G | A |
| PUDP | chrX | 7148220 | C | T |

**Section I: Robustness to Parameters**

Throughout our work we use a *p*-value threshold of 0.05 for filtering out samples with high expression correlation with immune cells according to Cibersort (Newman et al., 2015). However, increasing that threshold to 0.1 only removes one additional sample (**Figure S10a**). We use an FDR threshold of 0.1 (Benjamini–Yekutieli method) as a cutoff for statistical significance of differential ASE. This threshold is robust as the median number of significant genes changes by fewer than ten genes per sample for FDRs between 0.05 and 0.2 (**Figure S10b**). In our work we choose a threshold of 0.15 as a cutoff for a large effect size for differential ASE. The median number of genes called significant per sample changes roughly linearly with changes in this parameter at a fixed FDR of 0.1 (**Figure S10c**).

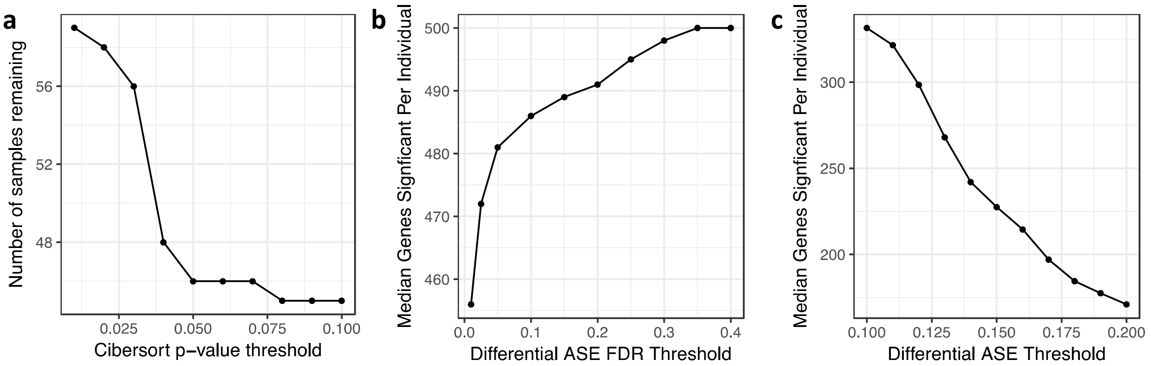


**Figure S10** Analysis is robust to parameters we selected. **a.** As we vary the Cibersort *p*-value threshold on the x-axis, the number of samples that remain in the study decreases, but only one more samples is removed between a threshold of 0.05 and 0.1. **b.** As we lower the FDR threshold for differential ASE (x-axis), fewer genes remain significant per individual. However, the median number of significant genes changes by fewer than ten genes per sample for FDRs between 0.05 and 0.2. **c.** We fix the FDR threshold to 0.1 and vary the threshold on differential ASE and find that the median number of genes significant per individual changes roughly linearly with the threshold.
